# A diffusion-driven switch specifies rhizoid precursor cells in *Marchantia polymorpha*

**DOI:** 10.1101/2023.04.04.535528

**Authors:** Josep Mercadal, Mar Ferreira-Guerra, Ana I. Caño-Delgado, Marta Ibañes

**Affiliations:** Departament de Física de la Matèria Condensada, Facultat de Física, Universitat de Barcelona, 08028 Barcelona, Spain; Universitat de Barcelona Institute of Complex Systems (UBICS), 08028 Barcelona, Spain; Department of Molecular Genetics, Centre for Research in Agricultural Genomics (CRAG), CSIC-IRTA-UAB-UB, Campus UAB (Cerdanyola del Vallès), 08193 Barcelona, Spain

**Author notes:** Max Planck Institute for Plant Breeding Research, 50829 Cologne, Germany (Josep Mercadal).

## Abstract

The specification of rhizoid precursor cells in the epidermis of *Marchantia polymorpha* gemmae has been shown to involve lateral inhibition mediated by the microRNA FRH1, which represses its activator RSL1, a rhizoid-specific transcription factor. However, how inhibition is conferred to adjacent cells and which is the mechanism underlying the emergence of rhizoid precursors remain unknown. In this paper, we use mathematical and computational modeling to show that the previously reported rhizoid patterns in WT, gain-of-function and lossof-function mutants of *FRH1* and *RSL1* are consistent with lateral inhibition mediated by a mobile FRH1. Our modeling results suggest that cells in *Marchantia* wildtype gemmae reside close to a critical state, where diffusion of FRH1 drives a switch of RSL1 expression that specifies rhizoid precursors. This process involves an initially random trigger and subsequent lateral inhibition, leading to cellular patterns consisting of small and filamentous clusters of rhizoid precursors. We confirm these predictions with new data on WT rhizoid distributions. Our findings highlight a novel mechanism of cellular pattern formation, opening new research directions for understanding cellular differentiation and tissue morphogenesis, with potential implications for a broad range of biological systems.

## Introduction

During organismal development, cells differentiate into distinct types to create specialized structures. This process typically involves universal mechanisms of pattern formation that reliably specify every particular cell type, tissue, and organ, with the precision and robustness so characteristic of biological forms. Several mechanisms are known to underlie cell-fate patterning, ranging from completely cell-autonomous to those relying on intercellular communication. For instance, the differentiation of photoreceptor cells in the eye of *Drosophila* is known to involve cell-autonomous random decisions caused by the stochastic expression of the *Spineless* gene [1, 2, 3]. In the sepals of *Arabidopsis thaliana*, fluctuations in the concentration of the transcription factor ATML1 generate a random pattern of giant cells interspersed between smaller epidermal cells [4]. In contrast, spatially periodic patterns can be achieved when there is cell-to-cell communication. For instance, cells interacting via lateral inhibition — where non cell-autonomous signals prevent adjacent cells from adopting the same fate — leads to characteristic salt-and-pepper patterns [5]. Lateral inhibition participates in many different cell specification processes in plants and animals, and can be mediated by distinct mechanisms. A paradigmatic example occurs within the Notch signaling pathway [6, 7, 8], where a transmembrane ligand (Delta) in a cell binds to the transmembrane receptors (Notch) of adjacent cells, triggering a signaling cascade that regulates cell differentiation [9, 10]. Lateral inhibition can also be mediated by repressive signals moving between adjacent cells, as happens in activator-inhibitor systems [11, 12]. If the activator is autocatalytic and does not diffuse, activator-inhibitor systems can exhibit patterns with two cell states, as occurs with the distribution of prestalk-prespore cell fates in the slime mould *Dyctiostelium discoideum* [13]. In the root epidermis of *Arabidopsis*, the specification of trichoblasts involves lateral inhibition through the movement of the MYB protein CAPRICE [14, 15], and competition for the hormone auxin mediates lateral inhibition in the specification of sieve elements and gap cells within the protophloem [16].

Recent research suggests that lateral inhibition plays a role in specifying rhizoid precursor cells in the epidermis of the common liverwort *Marchantia polymorpha* [17]. *M. polymorpha* has recently emerged as an exceptional model organism for studying the adaptations that allowed plants to transition from water to land [18, 19], thanks to its small genome, short life cycle, and ease of crossbreeding [20, 21]. Among the most decisive of these evolutionary novelties was the emergence of rooting systems [22, 23], which originally consisted of simple filamentous outgrowths called rhizoids, and were likely inherited from earlier adaptations in land plants’ algal ancestors [24].

In the gemma epidermis of *Marchantia*, rhizoid precursor cells are already specified on the first day after germination. This requires the activity of the bHLH transcription factor ROOT HAIR DEFECTIVE SIX-LIKE1 (MpRSL1) [25, 26]. MpRSL1 (from now on, RSL1) is known to induce the activity of the microRNA FEW RHIZOIDS1 (FRH1), which feeds back on RSL1 to repress its activity [17, 27]. These interactions constitute an activator-inhibitor system, with RSL1 acting as the activator and FRH1 as the inhibitor. The interplay between RSL1 and FRH1 controls the spatial distribution of rhizoid precursors, which consists of isolated cells and small clusters with linear or chain-like shapes (from now on *filamentous* clusters), indicating the presence of lateral inhibition preventing the formation of larger, more compact clusters [17]. Albeit FRH1 is required to mediate this lateral inhibition [17], the underlying molecular mechanisms are unknown. It has been suggested that FRH1 itself, by moving to adjacent cells, might be the main factor driving lateral inhibition [17, 25, 28]. The activator-inhibitor nature of this system could result in a Turing instability driving the rhizoid precursor fate; however, a detailed analysis testing this hypothesis has not been performed. By combining mathematical modeling and quantitative data analysis, we propose that rhizoid precursor patterns do not emerge from a Turing instability but through a diffusion-driven switch, where the adoption of distinct cellular fates crucially depends on the mobility of FRH1.

## Results

### Rhizoid specification is triggered cell-autonomously and randomly

In the gemmae of FRH1 loss-of-function mutants (*frh1*^*lo f*^), both epidermal and rhizoid precursor cells can be identified [17, 27], suggesting that the determination to develop into either cell type can occur without FRH1. Because in these mutants lateral inhibition is absent [17], we hypothesized that cell-fate decisions do not require intercellular crosstalk, and are therefore independent of adjacent cells. To test this premise, we compared published data of *frh1*^*lo f*^ mutants with a null model where every cell in a two-dimensional hexagonal array is allowed to become either a rhizoid precursor with probability *p*, or an epidermal cell with probability 1− *p* (Methods; Fig. 1A, B). We chose a value of *p* consistent with the average density of rhizoid precursors in *frh1*^*lo f*^ mutants, *p* = ⟨*ρ*⟩ ≈0.2 [28]. In this case, the patterns obtained with the null model predominantly consist of small clusters of rhizoid precursors and less frequent, larger clusters with more compact shapes (Fig. 1B, Left). These patterns are consistent with those observed in *frh1*^*lo f*^ mutants [17]. To quantitatively compare the phenotypes obtained using the null model with those of the mutants, we used two statistical metrics previously employed to quantify the properties of rhizoid precursor clusters [17]: the frequency of cluster sizes (*f*_*k*_; the proportion of clusters of size *k* among all clusters) and the neighbor density (*n*_*k*_; the average number of adjacent rhizoid precursors that a rhizoid precursor in a cluster of size *k* has; Methods). Our results show that the null model with *p* = 0.2 reproduces the distributions of *f*_*k*_ and *n*_*k*_ in *frh1*^*lo f*^ mutants (Fig. 1C, D), supporting the hypothesis that rhizoid specification in *frh1*^*lo f*^ mutants is random and cell-autonomous.

**Figure 1:**
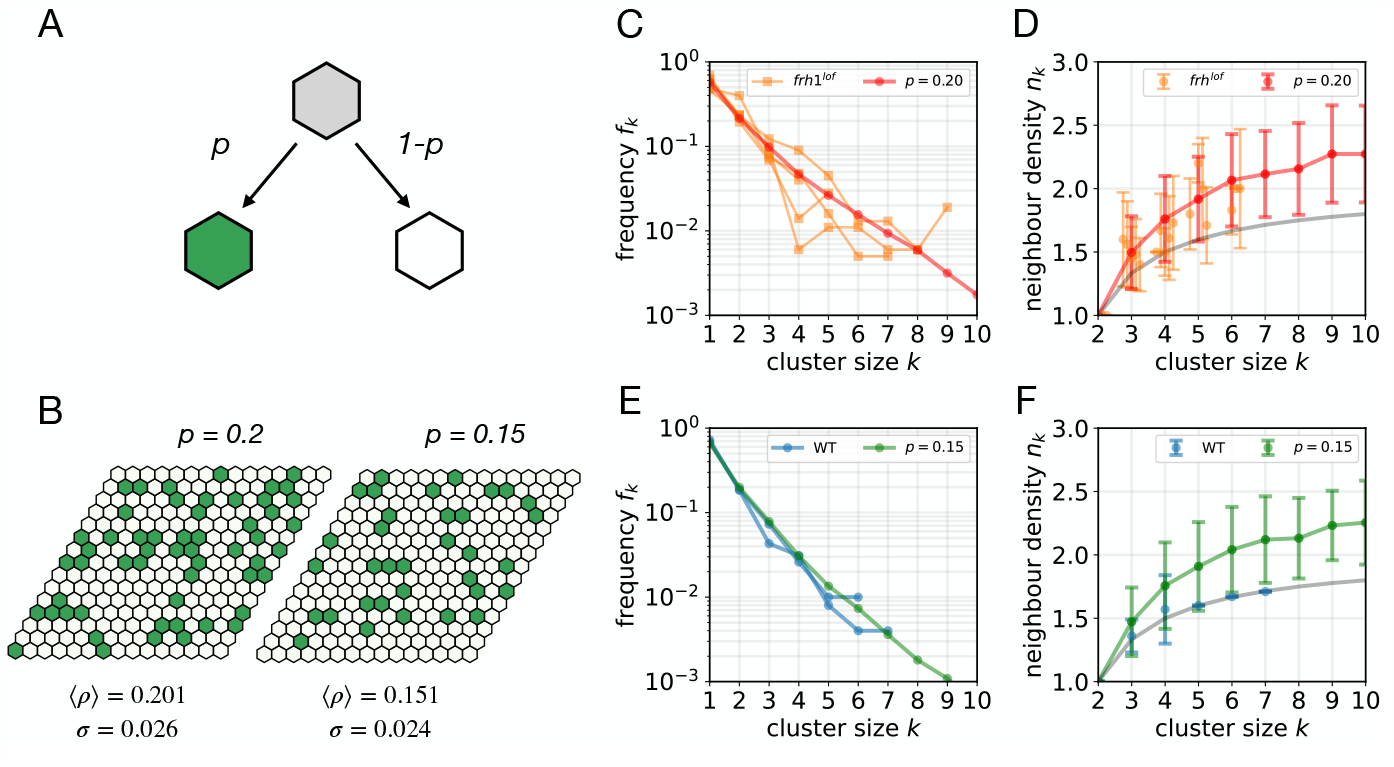
A cell-autonomous random model reproduces global, but not local properties of WT patterns. **A** In the null model, every cell can become either a rhizoid precursor (green) or a epidermal cell (white) with probability *p* and 1 *p*, respectively. **B** Typical patterns of rhizoid and epidermal cells in a lattice of 15×15 hexagonal cells, where every cell decision is independent from the other cells, for (Left) *p* = 0.2 and (Right) *p* = 0.15. ⟨*ρ*⟩ and *σ* denote, respectively, the mean and the standard deviation of the rhizoid density computed from *N* = 1000 lattices. **C, D** Results comparing the distribution of cluster sizes (*f*_*k*_, C) and the neighbor density (*n*_*k*_, D) of the null model for *p* = 0.2 (red) and four different FRH1 loss-of-function mutants (orange; data from [17]). **E, F** Same quantities as in C-D but comparing the null model for *p* = 0.15 (green) with Tak-1 and Tak-2 WT data [17] (blue). Points and error bars in D and F denote, respectively, the average value and ± the standard deviation of *n*_*k*_. The distribution of *n*_*k*_ is not symmetric, and cannot take lower values than those corresponding to filamentous clusters (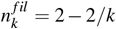; shown in gray). All the results are computed from *N* = 1000 simulations on hexagonal lattices of size 15 × 15 cells.

We next evaluated to what extent the mechanism of random, cell-autonomous binary decisions applies to WT patterns. To that end, we used the null model with a value of *p* consistent with the rhizoid precursor density in WT phenotypes (*p* = ⟨*ρ*⟩≈ 0.15 [17, 28]). In this case, most rhizoid precursors emerged either isolated or forming clusters of few cells (Fig. 1B, Right). For this value of *p*, the cluster size distribution *f*_*k*_ is consistent with its experimental counterpart (Fig. 1E, Supplementary Text S2.1). However, as expected, the values of *n*_*k*_ deviate significantly, most noticeably for larger clusters (Fig. 1F). This discrepancy is consistent with lateral inhibition acting on WT gemmae to turn rhizoid precursor clusters into more filamentous shapes [17], a feature absent in the null model. While the density ⟨*ρ*⟩ and *f*_*k*_ quantify global properties of the spatial distribution of rhizoid precursors, *n*_*k*_ is a measure of the local structure of clusters. Therefore, our results indicate that in WT gemmae, the global properties of rhizoid precursor patterns are consistent with a random, cell-autonomous binary decision, while the local structure of clusters is not.

### A mechanistic model for rhizoid precursor specification

Based on the previous results, we investigated the mechanistic basis of the random binary decisions underlying rhizoid precursor specification. Random binary cell fates are a well-known hallmark of bistable switches, provided sufficient variability in the initial conditions or stochasticity in the systems’ dynamics [29, 30, 31, 32, 33]. Because RSL1 is necessary for rhizoid formation [25, 26], we hypothesized that a cell-autonomous bistable switch of RSL1 activity might be involved in rhizoid specification. In WT gemmae, this switch will be modulated by FRH1, but not in *frh1*^*lo f*^ mutants. Mechanistically, bistable switches can be driven by positive feedback [29, 31, 34]. Accordingly, we assumed that RSL1 proteins (*R*) stimulate (directly or through intermediaries) the transcription of its own mRNA (*m*) in a positive feedback loop (Fig. 2A, Methods). Additionally, since FRH1 (*F*) is activated by RSL1, we assumed that RSL1 proteins induce the transcription of FRH1. Because FRH1 presumably inhibits RSL1 via mRNA cleavage [27], we set FRH1 to cleave RSL1 mRNA (Fig. 2A; Methods). We expect similar outcomes regardless of how we model this inhibition.

Using the levels of RSL1 proteins as a reporter of rhizoid precursor fate (high *R* for rhizoid precursors, low *R* for epidermal cells), we found that positive feedback of RSL1 can lead to bistability in the cell’s states: one with high levels of *m, R*, and *F* (rhizoid fate), and another with low levels (epidermal fate; Fig. S1). When the synthesis rate of FRH1 (represented by the parameter *γ*, or *inhibition strength*) exceeds a certain threshold, the resulting inhibition becomes too strong and disrupts the bistable switch. This disruption leads to the epidermal fate becoming the only stable state (Fig. S1A-C). FRH1’s requirement for lateral inhibition in WT rhizoid distributions [17] implies that it functions in a non-cell-autonomous manner. To include this regulation, we assumed FRH1 moves between cells in a diffusive manner, without any preferred direction and tending towards homogenization in the absence of reaction terms. With all of these assumptions, the mechanistic model constitutes an activator-inhibitor system in which the bistable nature of the activator is modulated by a mobile inhibitor that acts both cell-autonomously and in adjacent cells (Fig. 2A; Methods).

**Table 1:**
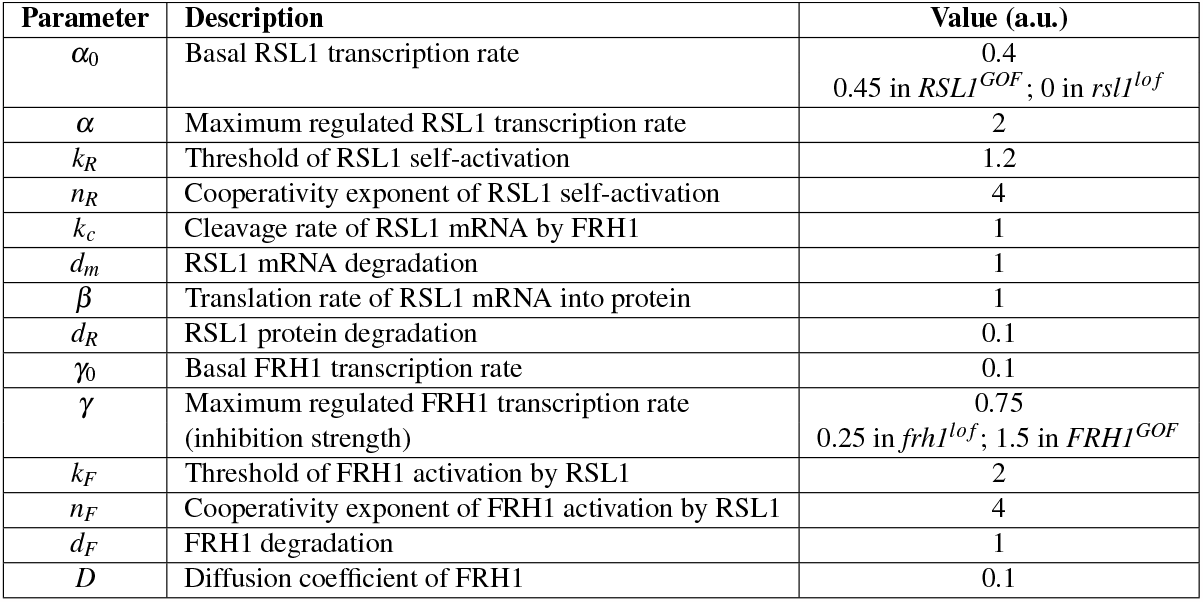
Parameters of the mechanistic model, their description and their default (WT) values (adimensional units). Parameter values used for the mutants are also specified. Default initial conditions are random **x**(*t* = 0) = **x**_0_𝒰 [0, 1]_15×15_ with **x**_0_ = (*m*_0_ = 0, *R*_0_ = 1.53, *F*_0_ = 0).

### The mechanistic model reproduces the experimental spatial distributions of rhizoid precursors in wild-type and in mutants

To test the validity of the mechanistic model, we studied its behavior on a two-dimensional regular hexagonal lattice, where every cell is mathematically described as a single point in space coupled to six adjacent neighbors (Methods). To incorporate the random component in cell-fate decisions, we assumed certain variability in the initial concentrations of RSL1 proteins. We explored different combinations of parameters and initial conditions, finding that the model could reproduce the experimental values of the density *ρ*, frequency distribution *f*_*k*_, and neighbor density *n*_*k*_ of clusters in WT gemmae (Fig. 2B-D, Table 1).

We next tested whether the model could also explain the experimental phenotypes of loss-of-function and gain-of-function mutants of FRH1 and RSL1 [17, 27]. To model these mutants, we assumed that the parameter values and initial conditions associated with the cells remain the same as in the WT, except for those parameters controlling transcription rates. To model the loss of function of FRH1, we decreased its transcription rate (from *γ* = 0.75 to *γ* = 0.25; Table 1). With this change, the mechanistic model reproduced the density ⟨*ρ*⟩, the distribution of cluster sizes *f*_*k*_, and the neighbor density *n*_*k*_ of *frh1*^*lo f*^ mutants (Fig. 2B; Fig. 2C,D Middle panels). Similarly, we modeled the gain of function of FRH1 by increasing its transcription rate (from *γ* = 0.75 to *γ* = 1.5; Table 1). Sufficiently large transcription of FRH1 results in patterns where all cells become epidermal, in agreement with the reported phenotypes of *FRH1*^*GOF*^ (Fig. S2). We applied the same procedure to model the gain and loss of function of RSL1. By setting the basal production of *m* to zero (*α*_0_ = 0; Table 1), we recovered the phenotypes of *rsl1*^*lo f*^ mutants, where no rhizoids appear (Fig. S2). Conversely, by increasing the basal production rate of RSL1 (from *α*_0_ = 0.4 to *α*_0_ = 0.45; Table 1) we observed that the resulting patterns reproduced the experimental statistics of ⟨*ρ*⟩, *f*_*k*_ and *n*_*k*_ of *RSL1*^*GOF*^ mutants (Fig. 2B; Fig. 2C,D Right panels). Hence, the mechanistic model captures the presence of long and compact clusters observed in *RSL1*^*GOF*^ mutants. Notably, despite the high activity of FRH1 in these mutants, the resulting patterns are consistent with the absence of lateral inhibition. Additionally, our results show that all these mutants’ phenotypes can also be reproduced by changes of *γ, γ*_0_ or *α*_0_ and of the initial conditions to other values (Fig. S3). Altogether, these results indicate that a mechanism where RSL1 and FRH1 interplay in a spatially extended activator-inhibitor system can explain the rhizoid precursor phenotypes observed in WT, *RSL1*^*GOF*^, *rsl1*^*lo f*^, *frh1*^*lo f*^ and *FRH1*^*GOF*^ mutants.

### An interplay between randomness and lateral inhibition

The activator-inhibitor nature of the mechanistic model can give rise to pattern formation via a Turing instability (Fig.3A; Methods). This instability occurs for specific parameter values, causing small perturbations to destabilize the homogeneous state. However, our analysis reveals that the rhizoid precursor patterns obtained in this region are not consistent with those seen in WT gemmae (Fig. S4). A detailed analysis of the mechanistic model indicates that WT phenotypes can only be reproduced in the parameter region close to a saddle-node bifurcation of the single-cell system (i.e. no diffusion), and only for specific initial conditions (Fig. 3A,B; Fig. S4). In this region, diffusion is necessary for rhizoid precursors to emerge (Fig. 3A,C). In the absence of diffusion, no rhizoid precursor is formed, while when FRH1 diffusion is large enough, rhizoid precursor patterns arise and resemble those observed in WT gemmae (Fig. 3C, S4).

**Figure 2:**
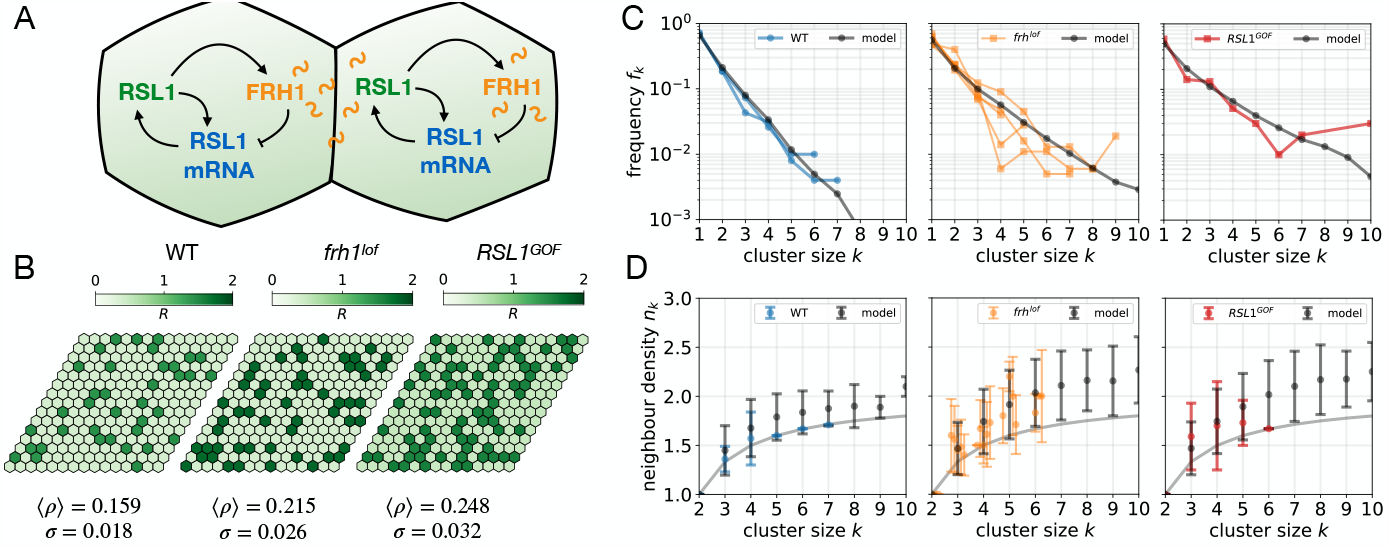
A mechanistic model for rhizoid patterns in wild-type and mutants. **A** Cartoon of the interactions between RSL1 and FRH1: RSL1 induces its own production through a positive feedback loop; RSL1 activates the production of FRH1, which in turn inhibits the translation of RSL1 into proteins by cleaving RSL1 mRNA, both cell and non-cell-autonomously. The non cell-autonomous action of FRH1 is mediated by the diffusion of FRH1 (wavy lines). **B** Examples of stationary rhizoid precursor patterns for WT (Left), *frh1*^*lo f*^ (Middle) and *RSL1*^*GOF*^ (Right), showing the activity of RSL1 (see Fig. S2 patterns of all the variables). ⟨*ρ*⟩ and *σ* denote, respectively, the average and standard deviation of the rhizoid precursor density. **C-D** Distribution of cluster sizes *f*_*k*_ (C) and neighbor density *n*_*k*_ (D) for WT (Left), *frh1*^*lo f*^ (Middle) and *RSL1*^*GOF*^ (Right). Experimental data from WT and mutants (color; data taken from [17]) is compared with simulations of the mechanistic model (black). For each genotype, the values of ⟨*ρ*⟩, *σ, f*_*k*_ and *n*_*k*_ in the model have been computed from *N*_*sim*_ = 1000 simulations using lattices of size 15 × 15. Parameter values of the model and initial conditions shown in Table 1.

**Figure 3:**
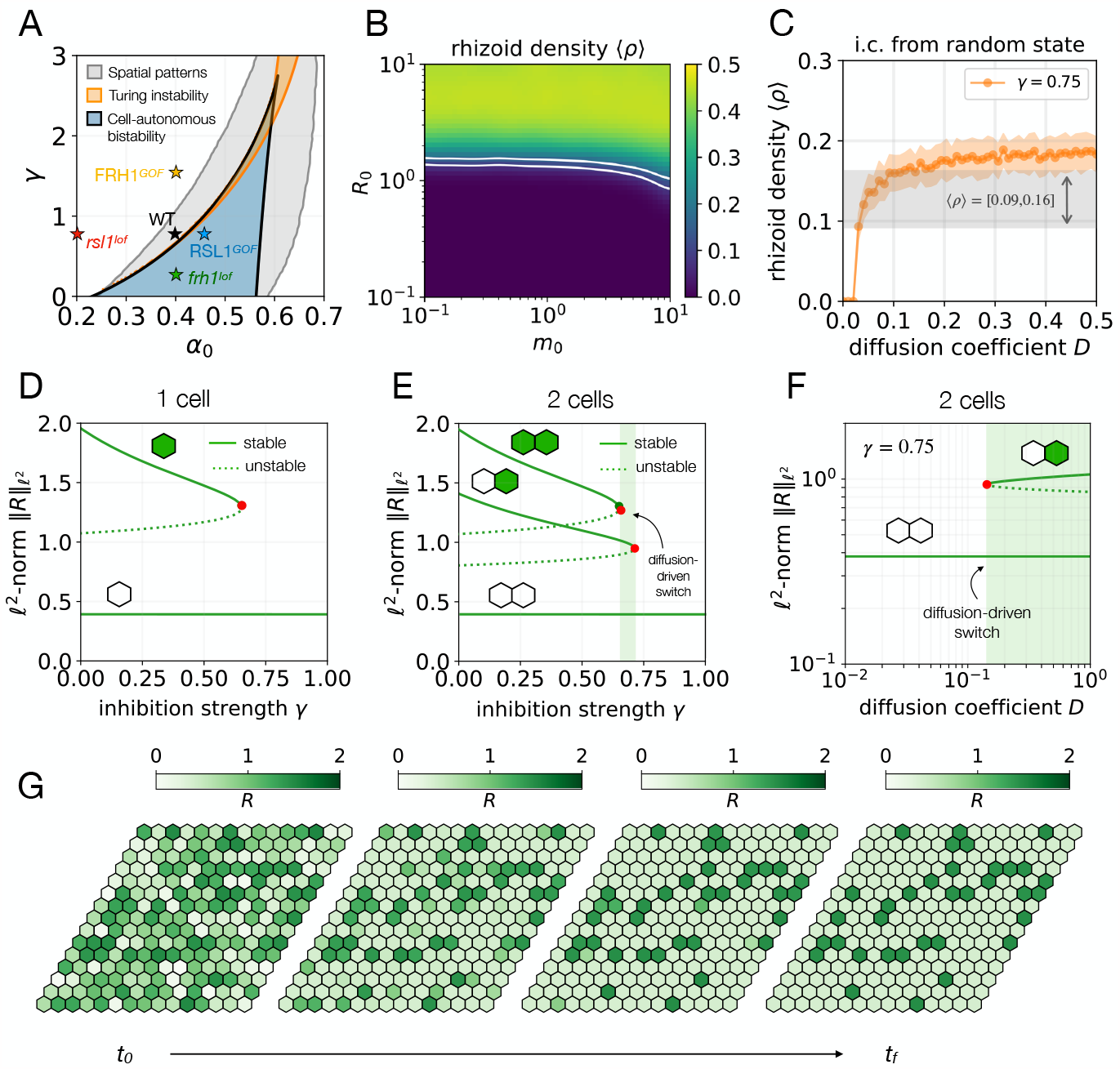
A diffusion-driven switch controls rhizoid patterning. **A** Stability diagram showing the possible steady-state regimes of the system in a 15 ×15 lattice, as a function of *γ* and *α*_0_ for FRH1 diffusion coefficient *D* = 0.1. Several regimes can be discerned: a region where patterns of rhizoid and epidermal cells can emerge (gray; it includes the blue and orange regions); a region of diffusion-driven linear instability (orange); a region of cell-autonomous (i.e. no diffusion) rhizoid/epidermal bistability (blue). **B** Average rhizoid density as a function of the amplitudes of the initial random variability in the levels of RSL1 (*R*_0_) and mRNA (*m*_0_), for *F*_0_ = 0. The two white lines enclose the region where this density is within the range of the WT, ⟨ *ρ*⟩∈ [0.09, 0.16]. Averages from *N* = 41 simulations with initial conditions **x**(*t* = 0) = **x**_*h𝒰*_ [0, 1]_15 ×15_, where **x**_*h*_ = (*m*_0_, *R*_0_, *F*_0_) and 𝒰 [0, 1]_15 ×15_ a uniform distribution of random numbers between 0 and 1 of size 15 15. **C** The emergence of rhizoid precursors depends on the diffusion coefficient *D*. For small values of *D*, rhizoid precursors cannot appear, because the only steady state of the system corresponds to the epidermal fate. For every value of *D*, circles represent averages of *ρ* over *N* = 41 simulations in a 15 ×15 hexagonal lattice. The orange shaded area denotes ± standard deviations. The gray-shaded area encloses the range of the WT density. **D-E** Bifurcation diagrams showing the steady states of the *ℓ*^2^-norm 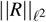 (Methods) for a single, isolated cell (D) and for two isolated cells coupled by diffusion of FRH1 (E). Red and green circles denote saddle-node and pitchfork bifurcations, respectively. The green-shaded area represents the region of diffusion-driven switch. The diffusion coefficient in E is *D* = 0.1. **F** Bifurcation diagram of the *ℓ*^2^-norm 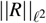 as a function of the diffusion coefficient *D* (with *γ* = 0.75), showing that the rhizoid precursor fate can not exist without diffusion. **G** Time evolution of typical rhizoid precursor patterns. For all panels, rest of the parameters as in Table 1. Initial conditions in C and G as in Table 1.

To further understand how rhizoid precursor patterns emerge in WT phenotypes, we analyzed the stationary states of a single-cell system and explored how these states were affected when introducing FRH1-mediated diffusive coupling between adjacent cells. Our results indicate that even when the inhibition strength is too high for cell-autonomous rhizoid specification, coupling two cells via diffusion of FRH1 can lead to one of them developing into a rhizoid precursor (Fig. 3D, E). For this to happen, the diffusion coefficient of FRH1 needs to surpass a certain threshold value, from which a new asymmetric state — the two cells acquiring opposite fates — emerges through a sharp switch (Fig. 3F). Moreover, higher diffusion enables rhizoid precursors to emerge for higher values of the inhibition strength (compare Fig. 3E with 3F). Our results also indicate that the larger the number of adjacent interacting cells, the smaller the diffusion coefficient needs to be for this switch to occur (compare Fig. 3C with 3F), and the larger the range of inhibition strengths capable of sustaining diffusion-driven patterns (Fig. S5). Overall, these findings suggest that diffusion of FRH1 generates a switch from a rhizoidless state to a pattern state where both rhizoid precursor and epidermal cells emerge.

Hence, when FRH1 does not diffuse, RSL1 is so strongly inhibited that the positive feedback cannot drive bistability in cell states (rhizoid precursor/epidermal cell). In contrast, when diffusion is present, RSL1 expression can switch to a high level and drive rhizoid precursor specification. The intuition behind this result is the following: when FRH1 diffuses, FRH1 inhibits less locally since a fraction of FRH1 moves to an adjacent cell. This drives stronger positive feedback of RSL1, enabling rhizoids to appear. Altogether, these results indicate that WT patterns emerge through a diffusion-driven switch, making the mobility of the microRNA FRH1 essential for rhizoid precursor specification in WT gemmae.

Additionally, because of the transport of FRH1 to adjacent cells, FRH1 diffusion also causes lateral inhibition. According to our analysis, a large diffusion coefficient of FRH1 can disrupt the local effect of lateral inhibition, even though the diffusion-driven switch still operates (Fig. S6). In such a case, rhizoid precursor patterns are similar to those obtained without lateral inhibition (Fig. S6). Based on these observations, we suggest that the range of FRH1 diffusion in *Marchantia* WT gemmae is not very large.

Our model also highlights the importance of initial cell-to-cell variability in establishing WT rhizoid precursor patterns. Because patterning happens through a switch, initial cell-to-cell variability must be sufficiently high to enable the emergence of rhizoid precursors (Fig. 3B, S4). Moreover, the initial average levels of RSL1 must be sufficiently low (Fig. 3B, S7). In this case, the inhibition mediated by FRH1 enables large clusters to arise and shapes them with filamentous shapes. Yet, because only some cells initially aim at reaching the rhizoid fate, the frequency of these large clusters is very small, whereas that of isolated rhizoid cells is much larger (Fig. 3G). Hence, to fully reproduce WT phenotypes in terms of rhizoid density, distribution of cluster sizes, and neighbor density, the model is required to use random initial conditions with sufficient variability between cells and low average RSL1.

On the basis of the models’ results in WT gemmae, we next evaluated how the diffusion-driven switch and lateral inhibition are modified in the gain-of-function and loss-of-function mutants. For *rsl1*^*lo f*^ and *FRH1*^*GOF*^ mutants, our model predicts the absence of rhizoid precursors, because of the lack of RSL1 in the former, and excessive inhibition in the latter. The switch is absent in both cases, because only the epidermal fate is stable (Fig. 3A; red and yellow stars). In contrast, in *frh1*^*lo f*^ mutants, the positive feedback of RSL1 is sufficiently strong to generate cell-autonomous bistable responses (Fig. 3A; green star). In this mutant, the combination of bistability and initial variability leads to cell-autonomous random decisions. In *RSL1*^*GOF*^ mutants, the RSL1 positive feedback, modulated by FRH1, is so strong that it can drive cell-autonomous bistability of epidermal/rhizoid fates, regardless of FRH1 diffusion (Fig. 3A; blue star). Thus, diffusion and thereby lateral inhibition become insignificant in *RSL1*^*GOF*^ and patterning is dominated by the cell-autonomous switch. Therefore, as in *frh1*^*lo f*^ mutants, the initial variability and the switch in *RSL1*^*GOF*^ mutants determines cell fate decisions independently of other cells.

Overall, these findings suggest that at the early stages of WT gemmae development, some cells might have higher values of RSL1, making them more likely to become rhizoid precursors. These cells start producing FRH1, which inhibits RSL1 cell-autonomously and in adjacent cells. This results in the inhibition of some high-RSL1 — turning them into epidermal cells — while others eventually differentiate into rhizoid precursors (Fig. 3G; Supplementary Movie 1). Taken together, the mechanistic model provides a dynamic description of rhizoid precursor specification in both WT and mutants.

### Lateral inhibition predominantly affects large clusters

The previous results pinpoint the importance of initially variability in generating rhizoid precursor patterns in WT gemmae. Analysis of the mechanistic model indicates that the initial random variability exhibited by cells confers robustness to changes in the inhibition strength (compare Fig. 4A-C to Fig. S7A-C). Furthermore, this variability reduces the sensitivity of the global pattern properties to the diffusion coefficient (Fig. S6). The random component of the initial conditions implies that lateral inhibition will not act uniformly on all cells at the outset. Specifically, because lateral inhibition of FRH1 regulates RSL1 activity, it can only affect cells that already have RSL1. This will have an impact on the effect of the inhibition strength, and therefore on how lateral inhibition influences the clusters of rhizoid precursors. For random initial conditions consistent with WT phenotypes, increasing the inhibition strength causes large clusters to become smaller and more filamentous, while having a relatively minor effect on small clusters (Fig. 4A). In contrast, when cells start with initially high RSL1 levels and low variability, increasing the inhibition strength does not lead to the same changes observed under WT initial conditions. Rather, the effect of lateral inhibition is similar regardless of the cluster size (Fig. S7).

**Figure 4:**
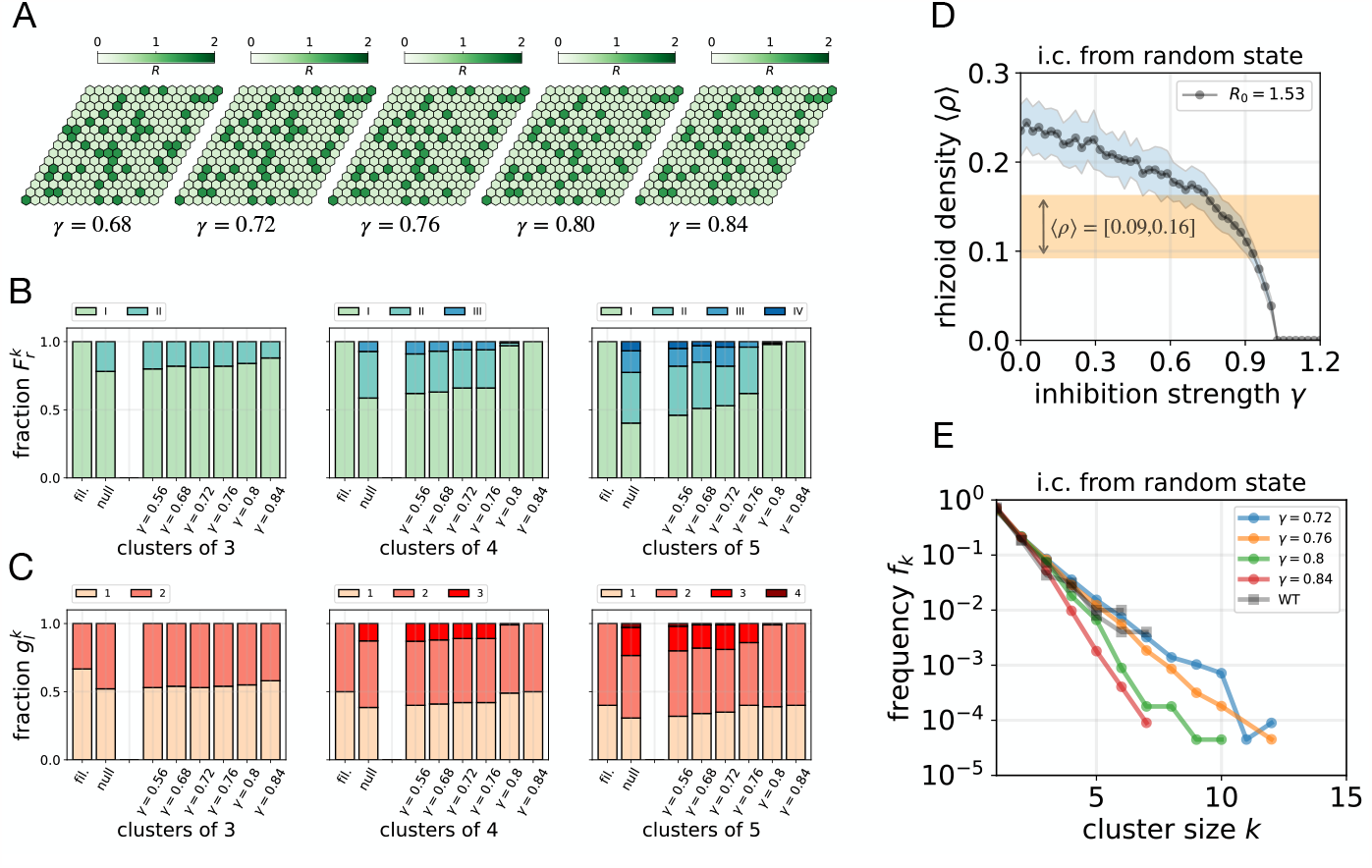
Lateral inhibition predominantly affects large clusters. **A** Examples of stationary patterns for different values of the inhibition strength *γ*. Initial conditions (*m*_0_ = *F*_0_ = 0, *R*_0_ = 1.53, as in Table 1) with the same seed generating the random numbers. **B** Frequency of cluster types 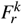 for different values of *γ*. The colors denote different cluster types (see Fig. S8). **C** Frequency of cells with *l* neighbors 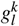 for different values of *γ*, for the same simulations as in B. Colors denote the number of adjacent rhizoid cells (*l* = 1, 2, 3, 4). For comparison, in B, C we also show the corresponding results for filamentous clusters (fil.) and for clusters obtained with the null model with *p* = 0.15. For each value of *γ* and each model, the results in B, C are computed with *N* = 1000 lattices of size 15 ×15. **D** Average rhizoid density as a function of *γ*. Points represent averages over *N* = 41 simulations, and shaded region ± their corresponding standard deviations. **E** Frequency distribution of cluster sizes for different values of *γ*, and compare to the WT of [17]. Same simulations as in B, C. In all panels, initial conditions are the same as in A.

The differential effect of lateral inhibition on distinct cluster sizes is also evidenced by comparing the neighbor density *n*_*k*_ of the mechanistic model in the WT with that of the null model (compare Fig. 1F with Fig. 2D). In both cases, random initial conditions and a switch-based decision lead to similar global properties of the pattern (⟨*ρ*⟩ and *f*_*k*_). However, in contrast to the null model, the lateral inhibition present in the mechanistic model has an impact on the shape of large clusters, turning them into more filamentous shapes.

To further quantify this differential effect of lateral inhibition, we classified all possible cluster configurations for every cluster size *k* in a hexagonal lattice (Fig. S8). There are *k* −1 ways of arranging cell clusters of size *k* based on the neighbor density *n*_*k*_ (Fig. S8; Methods). For instance, there is only one type of cluster of size *k* = 1 (isolated rhizoid cell), which has *n*_1_ = 0, and there is only one type of cluster of size *k* = 2, with *n*_2_ = 1. There are two types of clusters of size 3: filamentous (type *I*), with one rhizoid precursor in contact with two rhizoid precursors, each at one end of the filament 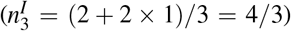; or compact (type *II*), with all three rhizoid precursors being adjacent to the other two 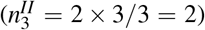. A similar reasoning applies to clusters of *k >* 3 (Fig. S8; Methods). For all cluster sizes, type I corresponds to filamentous clusters. To quantify the distribution of cluster types and their characteristics, we used the following two metrics: 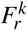, representing the frequency of clusters of size *k* and type *r* among all clusters of size *k*; and 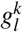, denoting the fraction of rhizoid precursors with *l* adjacent rhizoid precursors within all clusters of size *k* (*l* = 1, 2, …, *k*− 1). These metrics are not independent from each other for sizes *k* ≤4 (Supplementary Text S1). We computed 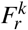 and 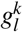 for the mechanistic model by counting all clusters within all simulated lattices. To assess the degree of randomness and lateral inhibition in the results of the mechanistic model, we compared the values of 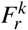 and 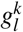 with those obtained in two limit cases: (1) the null model, i.e. a purely random model with no lateral inhibition, and (2) when all clusters are filamentous (type I). Because in WT gemmae large clusters appear only occasionally, we focused our analysis on sizes *k* = 3, 4, 5. For the patterns obtained with the mechanistic model, we found that clusters of sizes 4 and 5 become filamentous with smaller values of the inhibition strength *γ* than clusters of 3, which appear with the same proportions as in the null model (Fig. 4B, C). Taken together, our results predict that for the initial variability expected in the WT, the lateral inhibition involved in the diffusion-driven switch predominantly affects large clusters, decreasing the frequency of compact types and favouring filamentous shapes.

### New experimental data corroborate the predictions of the mechanistic model

We tested the predictions of our model by growing WT gemmae and visualizing rhizoid precursor cells through chloroplast autofluorescence (Fig. 5A; Methods). The frequency of cluster sizes *f*_*k*_ and neighbor density *n*_*k*_ were compatible with those previously reported [17], and were accurately reproduced by the mechanistic model (Fig. 5B, C). We observed large variability in the spatial distributions of rhizoid precursors, a feature also reproduced by the model (Fig. 5E). Additionally, we quantified the frequency of cluster types 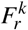 and the fraction of adjacent rhizoid precursors 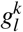 in all gemmae. Clusters of size *k* = 3 exhibited frequencies compatible with the mechanistic model (Fig. 5F,G). We classified clusters of size *k*≥ 4 into three groups: types I, II or higher than II. Analogously, we classified rhizoid precursors into those having one, two or more than two adjacent rhizoid precursors. The proportions 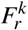 and 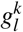 associated with this classification were statistically compatible with those of the mechanistic model, but not with those of the null model (Fig. 5F,G). As opposed to larger sizes, the statistics of size 3 clusters were also compatible with the null model, indicating that lateral inhibition is much less noticeable than in larger clusters (Fig. 5F,G). Hence, our quantitative data on rhizoid precursor distributions further confirm the predictions of the mechanistic model, supporting that rhizoid precursor patterns in WT gemmae emerge through a diffusion-driven switch and from an initially random pre-pattern of cellular states, where lateral inhibition influences rhizoid specification through the mobile microRNA FRH1.

**Figure 5:**
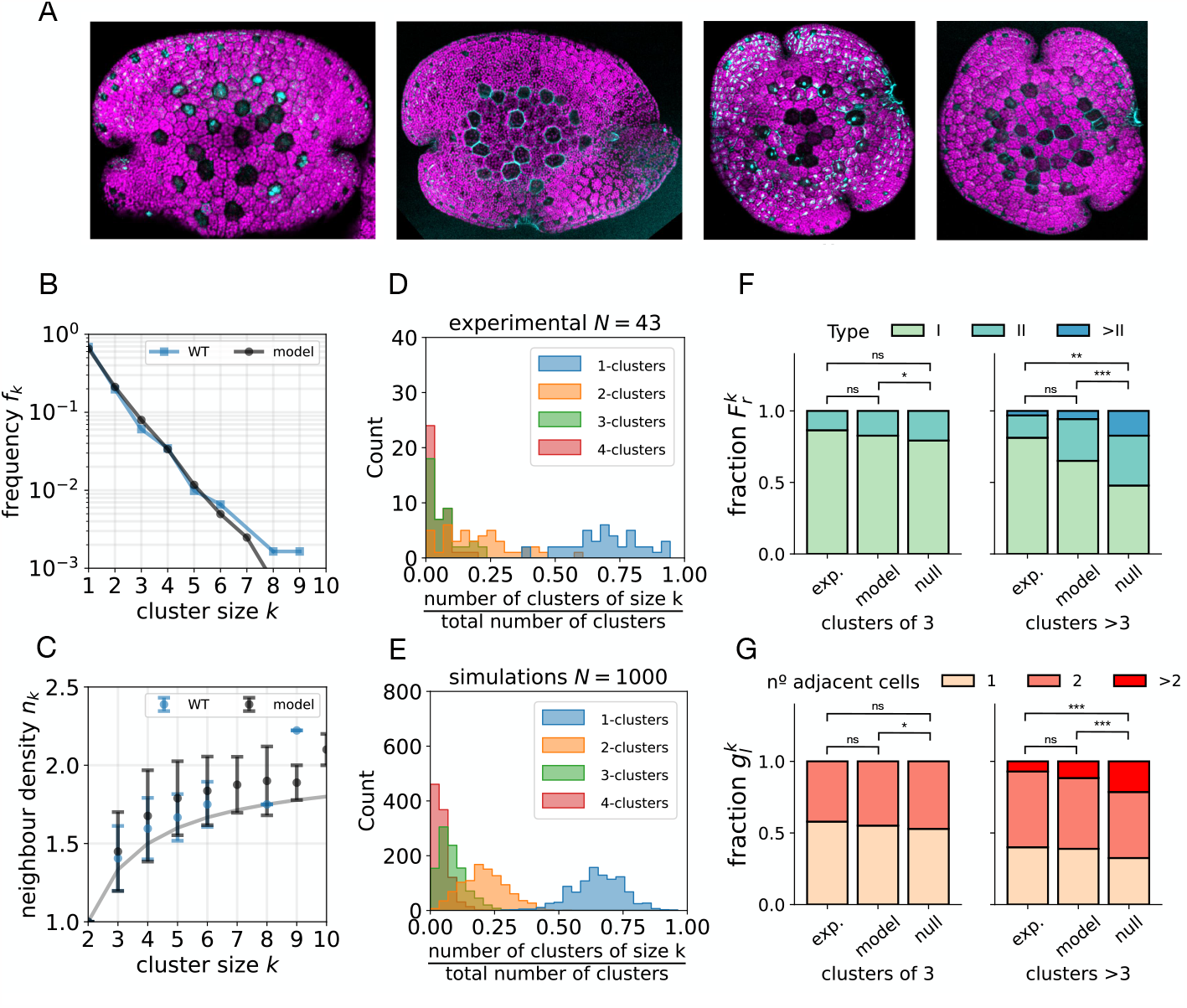
The results of the model are confirmed by new data on WT rhizoid distributions. **A** Four examples of the *Marchantia* WT gemmae used to compute the measures of spatial statistics (*N*_*gemmae*_ = 43). **B** Comparison between the distribution of cluster sizes *f*_*k*_ in WT gemmae (blue; *N*_*gemmae*_ = 43) and in that obtained with simulations of the mechanistic model (black; *N* = 1000, parameter values and initial conditions as in Table 1). **C** Comparison between the neighbor density of WT gemmae (blue) and simulations (black). **D-E** Histograms representing the frequency distribution of clusters within each gemmae, for the experiments (D) and for the simulations (E). **F** Side-by-side comparison of the fraction 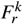 between experiments, the mechanistic model, and the null model with *p* = 0.15 (*N* = 1000). Due to the low number of large clusters, 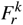 is separated in clusters of size *k* = 3 and *k >* 3. **G** Side-by-side comparison of the fraction 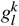 between experiments, the mechanistic model, and the null model. All the results of panels B-G come from the same simulations. ns *p*-value*>* 0.05; * *p*-values= 0.012, 0.021; ** *p*-value= 0.0013; \*\*\**p*-value*<* 0.0001. The broken down statistics and *p*-values in F and G can be found in Tables S5-S8.

## Discussion

Patterning processes are ubiquitous during the development of multicellular organisms. In this work, we propose that the specification of rhizoid precursor cells in the epidermis of *Marchantia polymorpha* gemmae is controlled by an activator-inhibitor system involving positive feedback of RSL1 and the mobility of the microRNA FRH1. Remarkably, our results indicate that the diffusion FRH1 is crucial for rhizoid specification in WT gemmae. In particular, diffusion facilitates a switch of RSL1 activity by decreasing the local inhibition of FRH1. In addition, the transport of FRH1 to adjacent cells confers lateral inhibition, turning rhizoid precursor clusters into filamentous shapes. In contrast to WT gemmae, the specification of rhizoid precursors in *f rh*1^*lo f*^ and *RSL*1^*GOF*^ are consistent with a cell-autonomous bistable switch. In these mutants, an initial, random pre-pattern determines the spatial distribution of rhizoid precursors.

The bistable switch of RSL1 activity proposed in our model is caused by a positive feedback loop. However, no experimental evidence is currently available supporting this specific mechanism. Even though we cannot discard other ways of driving a bistable switch, the overall consequences of bistability in RSL1 activity will be similar. Additionally, even though there is no direct evidence of the intercellular mobility of FRH1, previous studies have demonstrated the ability of other microRNAs to act non-cell-autonomously through intercellular movement [35, 36, 37, 38], thus supporting the plausibility of our assumption regarding the role of FRH1 mediating lateral inhibition.

Periodic spatial patterns in biological systems are commonly assumed to occur in a self-organized manner, initiated by the destabilization of spatially homogeneous states [10]. Paradigmatic examples of such processes are Turing patterns in reaction-diffusion systems [39, 40] and salt-and-pepper patterns driven by lateral inhibition mediated by cell-to-cell contacts [9, 8]. Interestingly, our mechanistic model displays a Turing instability, supporting the recently proposed notion that in discrete spatial lattices, a single morphogen is sufficient to generate periodic spatial patterning [41]. However, our results indicate that the spatial distribution of rhizoid precursors in WT gemmae is not consistent with a Turing instability, but it involves the interplay between initial random conditions and local lateral inhibition.

Turing and salt-and-pepper patterns emerging from linear instabilities of homogeneous states require long-range or local lateral inhibition, respectively [42]. In both cases, lossof-function mutants of genes mediating these inhibitions do not display patterned phenotypes. In *Marchantia* gemmae, rhizoid precursor patterns can emerge without lateral inhibition, as the phenotypes of *frh1*^*lo f*^ mutants demonstrate. Moreover, those in WT gemmae differ significantly from the salt-and-pepper patterns observed in conventional lateral inhibition mechanisms. In these cases, because lateral inhibition acts on all cells via cell-to-cell contacts, the proportion of inhibiting cells is much higher than those observed in rhizoid precursor patterns. Our modeling suggests that lateral inhibition in WT gemmae only acts in a fraction of cells and shapes the structure of clusters, but is not required for their emergence. Moreover, lateral inhibition is relatively weak, enabling clusters of few rhizoid precursors to persist.

In our model, the rhizoid precursor patterns of WT gemmae can only be reproduced in the parameter region close to the critical line separating cell-autonomous monostability and bistability. Although pattern formation requires non-zero diffusion — as in conventional reaction-diffusion systems — here it does so not through a linear instability of a homogeneous state but through a sharp switch, with a saddle-node bifurcation separating spatially homogeneous from spatially inhomogeneous ones. This *diffusion-driven switch* constitutes a novel mechanism for pattern formation in biological systems and can generate both random and periodic patterns depending on the initial states of the cells. Interestingly, cellular specification through a diffusion-driven switch can occur locally. This contrasts with cellular patterning driven by linear instabilities of homogeneous states, where cell specification occurs progressively through the propagation of differentiation fronts. Therefore, our results suggest that rhizoid precursors can become specified at different times during gemmae development, a process that might be especially relevant in regions of high proliferative activity, allowing the continuous generation of rhizoids during tissue growth.

The switch-like nature of rhizoid precursor specification suggests that pattern formation requires an initial pre-pattern of activity. This is most evident in *frh1*^*lo f*^ and *RSL1*^*GOF*^ mutants, where rhizoid precursors emerge cell-autonomously and randomly. Our results support that a random pre-pattern is necessary for initiating rhizoid specification in WT gemmae. Therefore, the final proportion of rhizoid precursors will be determined by the interplay between this initial prepattern and the lateral inhibition conferred by the diffusion of FRH1. Again, this contrasts with conventional pattern formation, where a self-organized mechanism — and not initial conditions — controls the proportion of cellular fates. Additionally, our analysis shows that initial variability in cellular states provides robustness to changes in the inhibition strength of FRH1. Our findings add to the growing body of knowledge on the prevalence of pre-patterning in biological systems, underscoring its relevance in pattern initiation and suggesting a role to enhance robustness [41].

Because the diffusion-driven switch can only occur close to a region of bistability, the underlying patterns are examples of *spatially localized states* — stable spatial structures embedded in a background of a different state [43, 44, 45]. Localized states appear in the so-called snaking region of the stability diagram — referring to the multiplicity of possible localized structures whose states can *snake* between different spatial configurations [46]. In the mechanistic model, the specification of rhizoid precursors in WT gemmae occurs in this region (Fig. 3A; gray region), where snaking phenomena of localized patterns can be observed for different parameter values (Fig. S5).

The proximity of WT patterns to a saddle-node bifurcation also suggests that rhizoid precursors reside close to a critical state. Criticality has long been proposed to endow biological systems with valuable features, including a balance between robustness against external perturbations and flexibility to adapt to changing environments, and granting optimal computational capabilities [47, 48]. In our model, criticality manifests in the long transients in the cell-autonomous dynamics before reaching the steady state (Fig. S1D, E). These *dynamical ghosts* [34] — referencing the steady-state remnant the system explores before collapsing to the true stable state — have been proposed to confer cells’ dynamic memory capabilities, enabling the processing and integration of time-varying signals [49]. Whether rhizoid precursors exploit this dynamic memory remains to be assessed, especially in light of the random nature of rhizoid specification.

Are the mechanisms of rhizoid specification conserved across early-diverging land plants? Do other closely related liverworts such as *Conocephalum conicum* or *Blasia pusilla* display similar rhizoid distributions? Rhizoid formation remains a poorly understood process, but the recently-disclosed mechanisms underlying their development in *Marchantia* pave the way to a hopefully burgeoning field, of which the combination of theory and experiment will be essential. Further investigation with new model organisms, phylogenetic reconstructions of early land plant lineages, and a theoretical understanding of the signaling pathways underlying their characteristic traits will shed light on the evolutionary significance of land colonization, and the molecular and physical processes that made such a remarkable transition possible.

We hypothesize that diffusion-driven switches underlie patterning processes not only in *Marchantia* but across a wide range of organisms, opening up new avenues for research in developmental and synthetic biology. Our findings highlight yet again the proclivity of living systems to operate close to a critical regime, prompting further investigation into how organisms utilize these states for adaptability, robustness, and survival.

## Methods

### Null model

To test how rhizoid precursors are distributed in *Marchantia* gemmae, we first use a *null model* where the specification is cell-autonomous, i.e. every cell performs a binary decision to randomly choose between becoming a rhizoid precursor or an epidermal cell. Accordingly, we model cell-fate decisions as Bernouilli trials, characterized by a single parameter, *p*, defined as the probability to become a rhizoid precursor (Fig. 1A). Conversely, *q* = 1 − *p* represents the probability to become an epidermal cell. Importantly, this model implies that all cells decide independently from the other cells. The null model is similar to previous model of rhizoid patterning in wild-type gemmae [17]. However, in contrast to that proposal, the null model differs in that cell decisions are independent of the states of any other cell, and does not include any intercellular interactions: no lateral inhibition nor any other rule that constraints where rhizoids can emerge based on where other rhizoids are placed. By fixing the size of the lattice (15× 15 cells) the statistics of the null model — including *f*_*k*_ and *n*_*k*_ — are fully determined by the value of the single, free parameter *p*. For each lattice, the rhizoid density, *ρ*, is defined as the fraction of cells that become rhizoid precursors. The total number of rhizoid precursors in a particular lattice follows a Binomial distribution of 15× 15 trials, each with probability *p*. The mean rhizoid density, ⟨*ρ*⟩, is computed by averaging the values of *ρ* over the total number lattices, i.e. 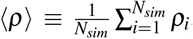, where the index *i* labels each simulation and *N*_*sim*_ is the total number of simulations. In the limit of an infinite lattice, this density satisfies *ρ*_∞_ = *p*. For smaller lattices, the density will only be similar to *p*, with deviations caused by the inherent fluctuations derived from random binary choices in a finite lattice.

### Mechanistic model

To model the dynamics of the RSL1-FRH1 system, we consider a regular hexagonal lattice where cells are labeled with the indices *i, j*, and assign the variables *m*_*i j*_, *R*_*i j*_, and *F*_*i j*_ to every cell. These variables represent the concentrations of RSL1 mRNA, RSL1 protein, and FRH1 microRNA, respectively. We model the rate of change of these concentrations with the following set of differential equations:

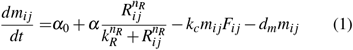

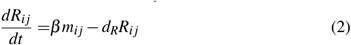

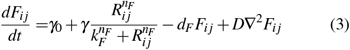

where ∇^2^ denotes a discrete Laplacian operator in a hexagonal lattice, i.e. ∇^2^*F*_*i j*_ = *F*_*i*+1, *j*−1_ + *F*_*i*+1, *j*_ + *F*_*i, j*−1_ + *F*_*i, j*+1_ + *F*_*i* − 1, *j*_ +*F*_*i* − 1, *j*+1_ − 6*F*_*i j*_. In this system, transcription of RSL1 mRNA happens with a constant rate *α*_0_ but is promoted by RSL1 proteins with maximum strength *α*, activation threshold *k*_*R*_, and cooperativity exponent *n*_*R*_. The degradation of RSL1 mRNA involves two contributions, one dependent on the FRH1 microRNA (cleaving) with rate *k*_*c*_, and another representing constant degradation of the mRNA by other means, with rate *d*_*m*_. The proteins RSL1 are translated from the mRNA at a constant rate *β* and are linearly degraded with rate *d*_*R*_. Transcription of FRH1 occurs with a basal rate *γ*_0_ and can be enhanced by the binding of RSL1 proteins to the FRH1 promoter, with a maximum rate *γ*, activation threshold *k*_*F*_, and cooperativity exponent *n*_*F*_. Finally, FRH1 can be linearly degraded with a constant rate *d*_*F*_, and can move from cell to cell with diffusion coefficient *D*. The strength of lateral inhibition performed by FRH1 depends on the cleavage rate *k*_*c*_ and on *γ*. In the main text, we refer to higher values of *γ* as stronger inhibition. Parameter values for WT, *frh1*^*lo f*^, *FRH1*^*GOF*^, *rsl1*^*GOF*^ and *RSL1*^*GOF*^ genotypes are shown in Table 1.

Studying the solutions of the system in the stationary state (i.e. all time derivatives set to zero) reveals that at the single-cell level — when *D* = 0 — the system can display bistability thanks to the self-activating nature of RSL1. While the low-RSL1 state is always stable, linear stability analysis reveals that the high-RSL1 state can lose stability in the presence of diffusion (Fig. 3A). Additionally, nonlinear inhomogeneous states exist close to the bistable region, which are not predicted by linear stability analysis (Fig. 3A; Fig. S5E).

### Lattice

To emulate the tissue layout of *Marchantia* gemmae, we performed all simulations on regular hexagonal lattices of size *N*_*x*_ ×*N*_*y*_ = 15 ×15, each hexagon representing a cell. This is consistent with the average number of cells enclosing the regions where rhizoid precursors appear [17, 28]. In the null model, no boundary conditions are provided because there are no cell interactions.

### Numerical simulations of the model

We solved the system of coupled differential equations (1-3) using the Runge-Kutta 4 method with a time step *dt* = 0.1. All the cluster statistics have been performed in the steady state, taken as the time *t* _*f*_ = 1000. We implemented reflecting boundary conditions, so that cells at the boundaries of the lattice only interact with their neighbors. For each variable (**x** ≡ (*m, R, F*)), initial conditions were drawn from a uniform distribution of random numbers fulfilling **x**(*t* = 0) = **x**_0𝒰_ [0, 1]_15×15_, where **x**_0_ ≡ (*m*_0_, *R*_0_, *F*_0_), and 𝒰 [0, 1]_15×15_ denotes a uniform distribution of 15× 15 (lattice size) random numbers between 0 and 1. We found that by setting *m*_0_ = *F*_0_ = 0 and only considering variability in RSL1 (*R*_0_ = 1.53) was sufficient. The same variability was used to model loss-of-function and gain-of-function mutants. For simulations where all cells start in the homogeneous state associated with the rhizoid precursor fate, we use a uniform distribution of random numbers of the form **x**(*t* = 0) = **x**_*h*_(1 +*V𝒰* [−1, 1]_15×15_), where here **x**_*h*_ are the values corresponding to the high-*R* homogeneous state. These initial conditions represent perturbations around this state with a variability parameter *V*. For simulations in the Turing region, *V* = 0.001 (Fig. S4B). Similar initial conditions were used when all cells started as rhizoid precursors but outside the Turing region (Fig. S4D,E). In this case, however, the values of **x**_*h*_ do not correspond to a stable homogeneous steady state, because out of this region the only stable homogeneous state is the one corresponding to all cells being epidermal. Therefore, in this region the density of the steady-state patterns will depend on the specific value of *V* .

The characteristic time *τ*_*c*_ associated with a ghost state (Fig. S1D) is defined as the time it takes for the system to reach the low-*R* steady state. To compute *τ*_*c*_, we start the system with initial conditions close to the proximal high-*R* steady state ((*m*_*h*_, *R*_*h*_, *F*_*h*_) ≈ (1.29, 1.29, 0.2); parameters as in Table 1) and let the system evolve until it reaches the true, low-*R* steady state ((*m*_*l*_, *R*_*l*_, *F*_*l*_) ≈ (0.38, 0.38, 0.1)). This has been performed for different values of the difference *ϕ* ≡ *γ* − *γ*_*c*_, where *γ*_*c*_ ≈ 0.665 is the bifurcation point. The characteristic times follow a power-law scaling of the form *τ*_*c*_ ∼ *ϕ*^*β*^, where the scaling exponent is *β* = −1*/*2 (Fig. S1E).

### Bifurcations and linear instabilities

The steady states of the mechanistic model and their stability have been computed via numerical continuation of saddle-node bifurcations with the Python package PyDSTool [50]. The cell-autonomous system displays bistability for certain parameter values (Fig. S1B,C; Fig. 3A, blue region). This bistable region emerges form a saddle-node bifurcation, where a pair of stable-unstable states are created in a discontinuous manner (Fig. S1B; red circles: saddle-node bifurcations). When coupled by the diffusion of FRH1, only the high-*R* steady state can become unstable through a linear instability (Fig. 3A; orange region). This can be shown by linear stability analysis of the system (Eqs. 1-3) [9, 51] (Supplementary Information). To quantify bistability and the emergence of inhomogeneous states, we use the *ℓ*^2^-norm of the variable *R*. In a hexagonal lattice, this quantity is defined as:

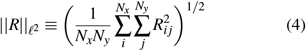

where *N*_*x*_ and *N*_*y*_ denote the total number of cells in each lattice direction. Intuitively, the *ℓ*^2^-norm quantifies the global state of the pattern (similar to the pattern density), and can be thought of as a measure of the energy associated with the spatial distribution of rhizoid precursors. In Fig. 3D-F we show this quantity as a function of *γ* and *D*. In Fig. S5B-D, ||*R*|| _*ℓ*_2 is shown for different number of cells. In both figures, saddle-node bifurcations are denoted with red circles, and green circles represent pitchfork bifurcations near the saddle nodes, which connect different stability branches through a series of ladders of asymmetric solutions (not shown) [52]. Thus, ||*R*|| _*ℓ*_2 provides a quantitative measure of the overall structure of the patterns, that can be used to characterize and make comparisons between them. The gray area shown in Fig. 3A encloses the region where ||*R*|| _*ℓ*_2 *>* 0, that is, where patterns are possible. For every pair (*γ, α*_0_), this region is computed by solving the system with high variability in initial conditions, *R*(*t* = 0) = *R*_0_(1 +*V𝒰* [−1, 1]_15×15_) (with *m*_0_ = *F*_0_ = 0 and *V* = 0.5), and by imposing the restriction ||*R*||_*ℓ*_2 *>* 0. Outside this region, no patterns are possible.

### Characterization of rhizoid precursor clusters

A cluster of size *k* corresponds to any spatial configuration of *k* rhizoid precursors that form a connected shape (i.e. every rhizoid precursor is adjacent to one or more rhizoid precursors) surrounded by epidermal cells. For instance, a cluster of size *k* = 1 is an isolated rhizoid precursor surrounded by epidermal cells. A cluster of size *k* = 2 consists of only two adjacent rhizoid cells completely surrounded by epidermal cells (Fig. S8). We only consider open clusters, i.e. those that do not form a closed loop.

The frequency of clusters of size *k, f*_*k*_, is defined as the total number of clusters of size *k* (*C*_*k*_) divided by the total number of clusters, obtained for all lattices:

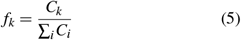

For each individual lattice, the ratio between the number of clusters of size *k* and the total number of clusters in that lattice can also be computed (Fig. 5E). The quantity *f*_*k*_ gives information about the global spatial organization of the pattern, but it does not inform about the local structure of clusters. For instance, one can imagine a lattice consisting exclusively of filamentous clusters, and another with all clusters being compact, but both of them with the same frequency of cluster sizes. Therefore, we used other metrics to characterize the local structure of clusters.

The neighbor density of clusters of size *k, n*_*k*_, is defined as the average proportion of adjacent rhizoid precursors that a rhizoid precursor has within a cluster of size *k*. Clusters of size *k* can be subdivided into *k* −1 different types, each defined by their value of *n*_*k*_ 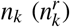 or, equivalently, by the total number of surrounding epidermal cells (Fig. S8). If *r* denotes the cluster type (*r* = *I, II*, …, *k* −1), then a closed-form expression for 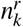 can be obtained for *k* ≤ 6:

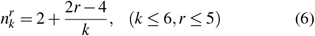

For every size *k*, the filamentous type (*r* = *I*) has the smallest possible value of *n*_*k*_, namely 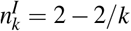. The most compact types (*r* = *k* −1) have the largest possible value of *n*_*k*_, which for *k*≤ 6 takes the form 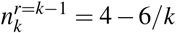. To obtain the mean neighbor density ⟨*n*_*k*_⟩ and the standard deviation for every size *k*, we use all the values of *n*_*k*_ obtained for each cluster from the total number of simulated lattices.

We denote as 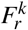 the proportion of clusters of size *k* and type *r* among all clusters of size *k* (from all simulated lattices). These proportions must sum up to one, and therefore the identity 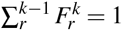 holds. The quantity 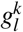 is the fraction of rhizoid cells within all clusters of size *k* (from all simulated lattices) that have *l* rhizoid precursor neighbors. The index *l* runs from 0 to *k* − 1 (for *k* = 1, 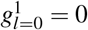). As before, these fractions sump up 1, 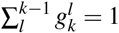.

The mean value of *n*_*k*_ depends on the proportion of clusters of each type, and therefore satisfies the relationships 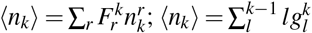, where summations run over all possible types and all possible number of rhizoid precursor neighbors, respectively. For *k* ≤ 3 the value of ⟨*n*_*k*_⟩ determines 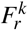 and 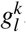, but this is not the case for *k* ≥ 4. Hence, the metrics ⟨*n*_*k*_⟩, 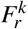, and 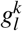 provide complementary information on the local structure of clusters. For clusters of size *k* = 1 and *k* = 2, ⟨*n*_*k*_⟩ and 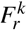 provide trivial information, as there is only one type of cluster. In contrast, for sizes *k >* 2 their values provide information on the mechanism driving rhizoid precursor specification, because the proportion of appearance of each cluster type will depend on the specific molecular mechanism at play. In Figs. 4,5 and Figs. S6,S7, these metrics are computed for the null model and for the mechanistic model for different parameter values and initial conditions. For the null model, these metrics can also be computed analytically for *k* ≤4 (Supplementary Text). Fig. 4 and Figs. S6,S7 also include these metrics when all clusters are filamentous (denoted as *f il*.). In this case, 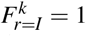 and 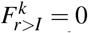 for any *k*, because all clusters are of the same type, and 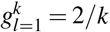 and 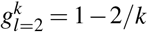 because all cells except the two at the borders (*k*− 2 cells in a cluster of *k* cells) have two adjacent rhizoid precursors, while the two cells at the borders have only one adjacent rhizoid precursor.

In Fig. 5F,G, the exact (two-sided) Fisher-test has been used to assess the statistical significance between the experimental results and the mechanistic and null models. To compute the statistical significance between the results of the mechanistic and null models, we used the Pearson *χ*^2^-test. The corresponding contingency tables can be found in the Supplementary Information (Tables S5-S8). All statistical tests have been computed with the *R* software.

### Numerical methods for cluster characterization

To compute the statistics of rhizoid precursors clusters, we combined custom-made software with the function scipy.ndimage.measurements.label from the Python library SciPy [53]. This function finds connected components of multi-dimensional arrays. In our case, we first identify the rhizoid precursors as the cells with *R*_*i j*_≥ 1. This threshold acts as an input for the label function to find connected components of rhizoid precursors. Each of these connected components constitutes a cluster, from which further information (*f*_*k*_, *n*_*k*_, 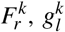) can be retrieved using custom-made methods.

### Experimental procedures

Propidium iodide (PI) staining to visualize cell walls was performed by incubating gemmae with 5 μg/ml PI for 10 minutes. Gemmae were subsequently washed twice with water and mounted on slides with heated 0.% agar solution. Imaging was performed after excitation by a Kr/Ar 488 nm laser using a FV 1000 confocal microscope (Olympus, Tokyo, Japan). PI and chlorophyll were detected with a band-pass 570-650 nm, and 650-750 nm filters, respectively. We counted all rhizoid precursor clusters, identifying their size and type, across all gemmae and computed the metrics *f*_*k*_, *n*_*k*_, 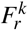 and 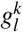 (without explicitly considering to which gemma each cluster belongs to).

## Acknowledgements

M.I. and J.M. acknowledge support from grant PID2021-125202NB-I00 funded by MCIN/AEI/10.13039/501100011033 and by “ERDF A way of making Europe”, and from the Generalitat de Catalunya through Grup de Recerca Consolidat 2017 SGR 1061. A.I.C-D. has received funding from the European Research Council (ERC) under the European Union’s Horizon 2020 research and innovation programme (grant agreement 683163). A.I.C-D. is a recipient of a BIO2020 grant funded by the Spanish Ministry of Economy and Competitiveness and Agencia Estatal de Investigación (MINECO/AEI) and Fondo Europeo de Desarrollo Regional (FEDER). J.M. acknowledges BES-2016-078218 funded by MCIN/AEI/10.13039/501100011033 and by “ESF Investing in your future”. M.F-G. is funded by the FPU fellowship (FPU16/06952) by the Ministry of Education, Culture and Sports.

## Author contributions

J.M. and M.I. designed the research with the help of M.F-G. and A.I.C-D. M.F-G. provided the experimental data. J.M. and M.I. formulated the mathematical models. J.M. analyzed the mathematical models and performed the numerical simulations. All authors analyzed the experimental data. J.M. and M.I. wrote the manuscript with the help of M.F-G. and A.I.C-D.

## Additional information

Additional information can be found in the Supplementary Text.

## References

[1] Mathias F Wernet, Esteban O Mazzoni, Arzu Ç elik, Dianne M Duncan, Ian Duncan, and Claude Desplan. Stochastic spineless expression creates the retinal mosaic for colour vision. Nature, 440(7081):174–180, 2006.

[2] Robert J. Johnston Jr.,, and Claude Desplan. Interchromosomal Communication Coordinates Intrinsically Stochastic Expression Between Alleles. Science, 343(6171):661–665, 2014.

[3] Michael Perry, Michiyo Kinoshita, Giuseppe Saldi, Lucy Huo, Kentaro Arikawa, and Claude Desplan. Molecular logic behind the three-way stochastic choices that expand butterfly colour vision. Nature, 535(7611):280–284, 2016.

[4] Heather M Meyer, José Teles, Pau Formosa-Jordan, Yassin Refahi, Rita San-Bento, Gwyneth Ingram, Henrik Jönsson, James C W Locke, and Adrienne H K Roeder. Fluctuations of the transcription factor ATML1 generate the pattern of giant cells in the Arabidopsis sepal. eLife, 6:e19131, 2017.

[5] Mark Eddison, Isabelle Le Roux, and Julian Lewis. Notch signaling in the development of the inner ear: Lessons from Drosophila. Proceedings of the National Academy of Sciences, 97(22):11692–11699, 2000.

[6] Spyros Artavanis-Tsakonas, Matthew D. Rand, and Robert J. Lake. Notch Signaling: Cell Fate Control and Signal Integration in Development. Science, 284(5415):770–776, 1999.

[7] Sarah J. Bray. Notch signalling in context. Nature Reviews Molecular Cell Biology, 17:722–735, 2016.

[8] Jelena Petrovic, Pau Formosa-Jordan, Juan C Luna-Escalante, Gina Abelló, Marta Ibañes, Joana Neves, and Fernando Giraldez. Ligand-dependent Notch signaling strength orchestrates lateral induction and lateral inhibition in the developing inner ear. Development, 141(11):2313–2324, 2014.

[9] Joanne R. Collier, Nicholas A. M. Monk, Philip K. Maini, and Julian H. Lewis. Pattern Formation by Lateral Inhibition with Feedback: a Mathematical Model of Delta-Notch Intercellular Signalling. J. Theor. Biol., 183:429–446, 1996.

[10] François Schweisguth and Francis Corson. Self-Organization in Pattern Formation. Developmental Cell, 49:659–677, 2019.

[11] Hans Meinhardt. Models of Biological Pattern Formation. Academic Press, 1982.

[12] Hans Meinhardt and Alfred Gierer. Applications of a Theory of Biological Pattern Formation Based on Lateral Inhibition. Journal of Cell Science, 15(2):321– 346, 1974.

[13] Hans Meinhardt. A model for the prestalk/prespore patterning in the slug of the slime mold Dictyostelium discoideum. Differentiation, 24(1-3):191–202, 1983.

[14] Natasha Saint Savage, Tom Walker, Yana Wieckowski, John Schiefelbein, Liam Dolan, and Nicholas A. M. Monk. A Mutual Support Mechanism through Intercellular Movement of CAPRICE and GLABRA3 Can Pattern the Arabidopsis Root Epidermis. PLoS Biology, 6(9):e235, 2008.

[15] S Schellmann, A Schnittger, V Kirik, T Wada, K Okada, A Beermann, J Thumfahrt, G Jürgens, and M Hülskamp. TRIPTYCHON and CAPRICE mediate lateral inhibition during trichome and root hair patterning in Arabidopsis. The EMBO Journal, 21(19):5036– 5046, 2002.

[16] Bernard Moret, Petra Marhava, Ana Cecilia Aliaga Fandino, Christian S. Hardtke, and Kirsten H. W. ten Tusscher. Local auxin competition explains fragmented differentiation patterns. Nature Communications, 11(1):2965, 2020.

[17] Anna Thamm, Timothy E. Saunders, and Liam Dolan. MpFEW RHIZOIDS1 miRNA-Mediated Lateral Inhibition Controls Rhizoid Cell Patterning in Marchantia polymorpha. Current Biology, 30:1905–1915, 2020.

[18] John L. Bowman. Insights into Land Plant Evolution Garnered from the Marchantia polymorpha Genome. Cell, 171(2):287–304, 2017.

[19] John L. Bowman. Walkabout on the long branches of plant evolution. Current Opinion in Plant Biology, 16:70–77, 2013.

[20] Kimitsune Ishizaki, Ryuichi Nishihama, Katsuyuki T Yamato, and Takayuki Kohchi. Molecular Genetic Tools and Techniques for Marchantia polymorpha Research. Plant and Cell Physiology, 57(2):262–270, 2016.

[21] Masaki Shimamura. Marchantia polymorpha: Taxonomy, Phylogeny and Morphology of a Model System. Plant Cell Physiol., 57(2):230–256, 2016.

[22] Richard M. Bateman, Peter R. Crane, William A. DiMichele, Paul R. Kenrick, Nick P. Rowe, Thomas Speck, and William E. Stein. Early Evolution of Land Plants: Phylogeny, Physiology, and Ecology of the Primary Terrestrial Radiation. Annu. Rev. Ecol. Syst., 29:263–292, 1992.

[23] Paul Kenrick and Peter R. Crane. The origin and early evolution of plants on land. Nature, 389:33–39, 1997.

[24] Paul Kenrick2 and Christine Strullu-Derrien. The Origin and Early Evolution of Roots. Plant Physiology, 166(2):570–580, 2014.

[25] Suvi Honkanen, Victor A. S. Jones, Giulia Morieri, Clement Champion, Alexander J. Hetherington, Steve Kelly, Hélène Proust, Denis Saint-Marcoux, Helen Prescott, and Liam Dolan. The Mechanism Forming the Cell Surface of Tip-Growing Rooting Cells Is Conserved among Land Plants. Current Biology, 26:3238–3244, 2016.

[26] Hélène Proust, Suvi Honkanen, Victor A.S. Jones, Giulia Morieri, Helen Prescott, Steve Kelly, Kimitsune Ishizaki, Takayuki Kohchi, and Liam Dolan. RSL Class I Genes Controlled the Development of Epidermal Structures in the Common Ancestor of Land Plants. Current Biology, 26:93–99, 2016.

[27] Suvi Honkanen, Anna Thamm, Mario A. Arteaga-Vazquez, and Liam Dolan. Negative regulation of conserved RSL class I bHLH transcription factors evolved independently among land plants. eLife, 7:e38529, 2018.

[28] Anna Thamm. The microRNA FEW RHIZOIDS1 controls rhizoid patterning in the liverwort Marchantia polymorpha. PhD Thesis, https://ora.ox.ac.uk/objects/uuid:5720a6a9-1640-484b-9709-ba56fc77f4b4, 2019.

[29] James E Ferrell Jr and Eric M Machleder. The biochemical basis of an all-or-none cell fate switch in xenopus oocytes. Science, 280(5365):895–898, 1998.

[30] Wen Xiong and James E Ferrell Jr. A positivefeedback-based bistable ‘memory module’that governs a cell fate decision. Nature, 426(6965):460–465, 2003.

[31] Attila Becskei, Bertrand Séraphin, and Luis Serrano. Positive feedback in eukaryotic gene networks: cell differentiation by graded to binary response conversion. The EMBO journal, 20(10):2528–2535, 2001.

[32] David Frigola, Laura Casanellas, José M Sancho, and Marta Ibañes. Asymmetric stochastic switching driven by intrinsic molecular noise. PloS one, 7(2):e31407, 2012.

[33] Arjun Raj and Alexander Van Oudenaarden. sNature, nurture, or chance: stochastic gene expression and its consequences. Cell, 135(2):216–226, 2008.

[34] Steven H. Strogatz. Nonlinear Dynamics and Chaos. CRC Press, 2014.

[35] Daniel H. Chitwood and Marja C. P. Timmermans. Small RNAs are on the move. Nature, 467:415–419, 2010.

[36] Daniel H. Chitwood, Fabio T.S. Nogueira, Miya D. Howell, Taiowa A. Montgomery, James C. Carrington, and Marja C.P. Timmermans.Pattern formation via small RNA mobility. Development, 23:549–554, 2009.

[37] Damianos S. Skopelitis, Kristine Hill, Simon Klesen, Cristina F. Marco, Patrick von Born, Daniel H. Chitwood, and Marja C. P. Timmermans. Gating of miRNA movement at defined cell-cell interfaces governs their impact as positional signals. Nature Communications, 9(3107), 2018.

[38] Liu Liu and Xuemei Chen. Intercellular and systemic trafficking of RNAs in plants. Nature Plants, 4:869– 878, 2018.

[39] Alan M. Turing. The chemical basis of morphogenesis. Phil Trans B, 237:37–72, 1952.

[40] Shigeru Kondo and Takashi Miura. Reaction-Diffusion Model as a Framework for Understanding Biological Pattern Formation. Science, 329(5999):1616–1620, 2010.

[41] Sheng Wang, Jordi Garcia-Ojalvo, and Michael B Elowitz. Periodic spatial patterning with a single morphogen. Cell Systems, 13(12):1033–1047.e7, 2022.

[42] Hans Meinhardt and Alfred Gierer. Pattern formation by local self-activation and lateral inhibition. BioEssays, 22:753–760, 2000.

[43] Edgar Knobloch. Spatially localized structures in dissipative systems: open problems. Nonlinearity, 21(4):T45–T60, 2008.

[44] Alan R. Champneys, Fahad Al Saadi, Victor F. Verônica A. Grieneisen, Athanasius F.M.Mareé, Nicolas Verschueren, and Bert Wuyts. Bistability, wave pinning and localisation in natural reaction–diffusion systems. Physica D, 416:132735, 2021.

[45] Victor Breña-Medina and Alan Champneys. Subcritical Turing bifurcation and the morphogenesis of localized patterns. Physical Review E, 90(3):032923, 2014.

[46] John Burke and Edgar Knobloch. Homoclinic snaking: structure and stability. Chaos, 17(3):037102, 2007.

[47] Thierry Mora and William Bialek. Are Biological Systems Poised at Criticality? Journal of Statistical Physics, 144:268–302, 2011.

[48] Miguel A. Muñoz. Colloquium: Criticality and dynamical scaling in living systems. Reviews of Modern Physics, 90:031001, 2018.

[49] Angel Stanoev, Akhilesh P Nandan, and Aneta Koseska. Organization at criticality enables processing of time-varying signals by receptor networks. Molecular Systems Biology, 16:e8870, 2020.

[50] Clewley RH, Sherwood WE, LaMar MD, and Guckenheimer JM. PyDSTool, a software environment for dynamical systems modeling. http://pydstool.sourceforge.net, 2007.

[51] M. C. Cross and P. C. Hohenberg. Pattern formation outside of equilibrium. Rev Mod Phys, 64(3):851, 2010.

[52] John Burke and Edgar Knobloch. Snakes and ladders: Localized states in the Swift–Hohenberg equation. Physics Letters A, 360(6):681–688, 2007.

[53] Pauli Virtanen and SciPy 1.0 Contributors. SciPy 1.0: fundamental algorithms for scientific computing in Python. Nature Methods, 17(13):261–272, 2020.

