## Supplementary Information for "A diffusion-driven switch specifies rhizoid precursor cells in *Marchantia polymorpha*"

#### 1 Relationship between $F_r^k$ and $g_l^k$ for $k \leq 4$

For every size $k$, there are $k-1$ values of $F_r^k$ and $g_l^k$. The latter can be obtained directly from the former for $k \leq 4$ (they constitute an algebraic system of $k-1$ linear equations with $k-1$ variables, for $k \leq 4$). This is because for a given size $k \leq 4$, all cluster configurations of the same type $r$ have same number of rhizoid precursors with $l$ adjacent rhizoid precursors (Fig. S8). Hence, $F_r^k$ and $g_l^k$ are not independent metrics for $k \leq 4$. We next exemplify this for $k = 3$ first. The fraction of rhizoid precursors within clusters of size $k = 3$ with $l = 1$ and with $l = 2$ adjacent rhizoid precursors is given by:

$$g_{l=1}^{k=3} = \frac{2}{3} F_{r=I}^{k=3}, \qquad g_{l=2}^{k=3} = \frac{1}{3} F_{r=I}^{k=3} + F_{r=II}^{k=3} \tag{1}$$

where the fractions $2/3$ and $1/3$ represent the proportions of rhizoid precursors with one or two neighbours, respectively, that appear with frequency $F_{r=I}^{k=3}$. Because clusters of type II have all cells with two neighbours, the factor in front of $F_{II}^3$ is just 1. Because these proportions must sum up to unity, the identity $\sum_{l=1,2} g_1^{k=3} = F_{r=I}^{k=3} + F_{r=II}^{k=3} = 1$ holds.

For clusters of size $k = 4$, the fractions of rhizoid precursors with $l$ rhizoid neighbors for $l = 1,2,3$ are:

$$g_{l=1}^{k=4} = \frac{1}{2} F_I^{k=4} + \frac{2}{4} F_{II}^{k=4}, \quad g_{l=2}^{k=4} = \frac{1}{2} F_I^{k=4} + \frac{1}{2} F_{II}^{k=4} + \frac{1}{2} F_{III}^{k=4}, \quad g_{III}^{k=4} = \frac{1}{4} F_{II}^{k=4} + \frac{1}{2} F_{III}^{k=4} \tag{2}$$

As before, we have the identity $\sum_{l=1,2,3} g_l^{k=4} = \sum_{r=I,II,III} F_r^{k=4} = 1$. These relationships are independent of the model underlying rhizoid formation.

For $k > 4$, each cluster type, which is defined by a value of $n_k$, can have different structures/shapes each involving different fractions of cells with $l$ adjacent rhizoids. Hence, $F_k^r$ and $g_l^k$ are not equivalent metrics for $k > 4$.

#### 2 Analytical expressions for the spatial metrics in the null model

We can obtain analytical formulas for the metrics of cluster statistics of the null model. In the null model, the probability of finding a specific cluster configuration of size $k$ is $p^k(1-p)^{e_k^r}$, where $e_k^r$ is the number of epidermal cells surrounding the cluster. For instance, for a cluster of rhizoid precursors of size $k = 3$ forming a

*Present address: Max Planck Institute for Plant Breeding Research, 50829 Cologne, Germany

horizontal, filamentous line, this probability is $p^3(1-p)^{10}$, because the cluster consists of 3 rhizoid precursors — each one appearing with probability $p$ — surrounded by 10 epidermal cells — each appearing with probability $1-p$. Because all configurations of the same type $r$ and size $k$ have the same value $e_k^r$ (Methods; Fig. S8), the probability for them to appear in the null model is the same. Therefore, the proportion of clusters of size $k$ and type $r$ in the null model is:

$$P_r^k = S_r^k p^k (1-p)^{e_k^r} \tag{3}$$

where $S_r^k$ is the total number of spatial configurations of clusters of size $k$ and type $r$ (Fig. S8). For instance, for clusters of size $k=3$ and type I, $S_I^3 = 9$ (Fig. S8). The probability for a cluster of size $k$ to appear, regardless of the type, is:

$$P_k = \sum_{r=I}^{k-1} P_r^k = \sum_{r=I}^{k-1} S_r^k p^k (1-p)^{e_k^r} \tag{4}$$

where the summation runs over all possible cluster types of size $k$. For $k=3$ there are only 2 types, I and II (Fig. S8). Type II is characterized by $e_3^{II} = 9$ surrounding epidermal cells, and it has two possible configurations, $S_{II}^3 = 2$ (Fig. S8). Hence, $P_3 = 9p^3(1-p)^{10} + 2p^3(1-p)^9$. Applying this to clusters of size $k < 5$, we find:

| $k$ | 1 | 2 | 3 | | 4 | | |
|---|---|---|---|---|---|---|---|
| $r$ | I | I | I | II | I | II | III |
| $e_k^r$ | 6 | 8 | 10 | 9 | 12 | 11 | 10 |
| $S_k^r$ | 1 | 3 | 9 | 2 | 29 | 12 | 3 |
| $P_k^r$ | $p(1-p)^6$ | $3p^2(1-p)^8$ | $9p^3(1-p)^{10}$ | $2p^3(1-p)^9$ | $29p^4(1-p)^{12}$ | $12p^4(1-p)^{11}$ | $3p^4(1-p)^{10}$ |

**Table S1:** Quantities associated with different cluster configurations, together with their associated probabilities of appearance in the null model.

With these probabilities we can determine analytical expressions for $f_k$, $n_k$, $F_r^k$ and $g_l^k$.

#### 2.1 Frequency of cluster sizes $f_k$ in the null model

The frequency of clusters of size $k$ among the total number of clusters is defined as:

$$f_k = \frac{P_k}{\sum_{k=1,2,\dots} P_k} \tag{5}$$

The identity $\sum_{k=1,2,\dots} f_k = 1$ holds. For the values of $p$ observed in WT gemmae, most clusters are of size $k \leq 4$, and therefore $P_{k>4} \ll P_4$. Then we can approximate

$$\sum_{k=1,2,\dots} P_k \approx P_1 + P_2 + P_3 + P_4 \tag{6}$$

and obtain closed formulas for $f_k$ in terms of $p$ using the expressions in Table S1. For $p = 0.15$, these formulas result in $f_1 = 0.6685$, $f_2 = 0.2141$, $f_3 = 0.0878$, $f_4 = 0.0396$ which coincide with the simulation results of the null model for $N = 1000$ lattices (Fig. 1), validating the analytical and the numerical approaches. Moreover, these values are in agreement with those of WT gemmae [1] (Fig. 1). Hence, the null model, with its only free parameter, $p$, chosen to be the average density of rhizoid precursors in WT gemmae ($p = 0.15$), reproduces both the density and frequency of cluster sizes of WT rhizoid patterns.

#### 2.2 Frequency of clusters of size $k$ and type $r$ ($F_r^k$) in the null model

The frequency of clusters of type $r$ among all clusters of a certain size $k$ is defined as:

$$F_r^k = \frac{P_r^k}{P_k} \tag{7}$$

with $\sum_{r=I,\ldots,k-1} F_r^k = 1$. From Table S1, we find the following closed-form expressions for $k \leq 4$:

$$F_{r=I}^{k=1} = 1 \quad (8)$$

$$F_{r=I}^{k=2} = 1 \quad (9)$$

$$F_{r=I}^{k=3} = \frac{9p^3(1-p)^{10}}{9p^3(1-p)^{10} + 2p^3(1-p)^9}, \quad F_{r=II}^{k=3} = 1 - F_{r=I}^{k=3} \quad (10)$$

$$F_{r=I}^{k=4} = \frac{29p^4(1-p)^{12}}{29p^4(1-p)^{12} + 12p^4(1-p)^{11} + 3p^4(1-p)^{10}}, \quad (11)$$

$$F_{r=II}^{k=4} = \frac{12p^4(1-p)^{11}}{29p^4(1-p)^{12} + 12p^4(1-p)^{11} + 3p^4(1-p)^{10}}, \quad F_{r=III}^{k=4} = 1 - F_{r=I}^{k=4} - F_{r=II}^{k=4} \quad (12)$$

These formulas can be substituted into Equations 1 and 2 to obtain analytical expressions for the fractions $g_l^k$ of the null model. By using different values of $p$, we compared these analytical expressions with the quantities $F_r^k$ and $g_l^k$ obtained from $N = 1000$ simulations of the null model, obtaining good agreement and validating the computational approach to extract and characterize clusters.

### 3 Linear stability analysis of the mechanistic model

Our mechanistic model consists on the following system of coupled differential equations:

$$\frac{dm_{ij}}{dt} = \alpha_0 + \alpha \frac{R_{ij}^{n_R}}{k_R^{n_R} + R_{ij}^{n_R}} - k_c m_{ij} F_{ij} - d_m m_{ij} \quad (13)$$

$$\frac{dR_{ij}}{dt} = \beta m_{ij} - d_R R_{ij} \quad (14)$$

$$\frac{dF_{ij}}{dt} = \gamma_0 + \gamma \frac{R_{ij}^{n_F}}{k_F^{n_F} + R_{ij}^{n_F}} - d_F F_{ij} + D\nabla^2 F_{ij} \quad (15)$$

where $\nabla^2 F_{ij} = F_{i+1,j-1} + F_{i+1,j} + F_{i,j-1} + F_{i,j+1} + F_{i-1,j} + F_{i-1,j+1} - 6F_{ij}$. To find the stability of the high-$R$ homogeneous state, we first numerically compute the fixed points of the uncoupled, cell-autonomous system. With these values, we consider a perturbation of the form $\delta \mathbf{x}_{ij} = \mathbf{x}_{ij} - \mathbf{x}_h$, where $\mathbf{x}_{ij} \equiv (m_{ij}, R_{ij}, F_{ij})$ is a vector of all the system's variables and $\mathbf{x}_h \equiv (m_h, R_h, F_h)$ denotes the values of these variables in the homogeneous steady state. We assume perturbations of the form:

$$\delta \mathbf{x}_{jk} = \sum_p^{N_x} \sum_q^{N_y} \mathbf{A}_{pq} e^{2\pi i(\frac{2\pi jp}{N_x} + \frac{2\pi kq}{N_y})} e^{\sigma_{pq} t} \quad (16)$$

where we switched notation from the dummy indices $i,j$ to $j,k$ to avoid confusion with the imaginary unit $i = \sqrt{-1}$. The homogeneous state will become unstable when $\mathrm{Re}(\sigma_{pq}) > 0$. In terms of the system's parameters, this condition can be found by computing the eigenvalues of the Jacobian matrix:

$$\mathbb{J} = \begin{pmatrix} -k_c F_h - d_m & \alpha n_R \frac{k_R^{n_R} R_h^{n_R - 1}}{(k_R^{n_R} + R_h^{n_R})^2} & -k_c m_h \\ \beta & -d_R & 0 \\ 0 & \gamma n_F \frac{k_F^{n_F} R_h^{n_F - 1}}{(k_F^{n_F} + R_h^{n_F})^2} & -d_F - 6D + D\Omega_{pq} \end{pmatrix} \quad (17)$$

where $\Omega_{pq} \equiv 2\cos\left(\frac{2\pi p}{N_x} - \frac{2\pi q}{N_y}\right) + \cos\left(\frac{2\pi p}{N_x}\right) + \cos\left(\frac{2\pi q}{N_y}\right)$, and $p,q$ are the Fourier modes associated with the non-homogeneous perturbations. The eigenvalues of $\mathbb{J}$ are the growth rates $\sigma_{pq}$, and therefore the sign of these eigenvalues will dictate the stability of the homogeneous state. In our case, the eigenvalue equation takes the form of a rank-3 polynomial $\lambda^3 + A\lambda^2 + B\lambda + C = 0$, where the coefficients $A,B,C$ are constants associated to the Jacobian $\mathbb{J}$. This equation can be solved numerically for different parameter values. In particular, in Fig. 3A we show the condition $\mathrm{Re}(\lambda) > 0$ in the plane $(\alpha_0, \gamma)$ (orange region).

### Supplementary Figures

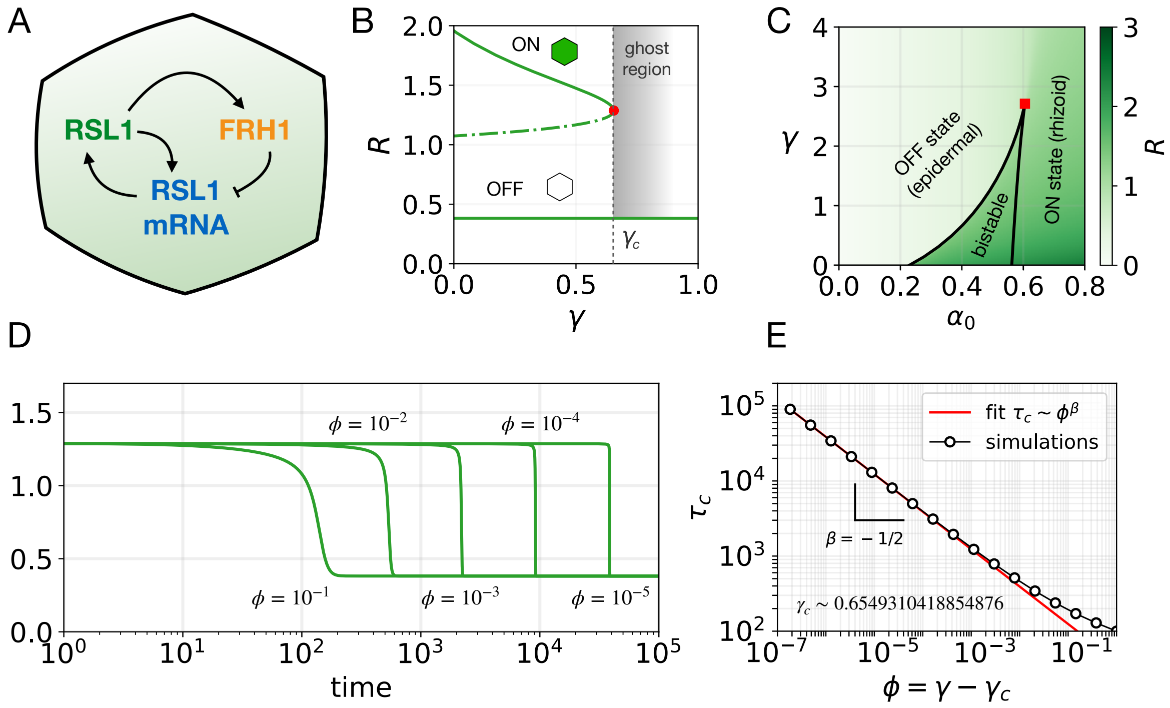


**Figure S1: Strong lateral inhibition by FRH1 can destroy the bistable switch.** **A** Sketch of the interactions of the mechanistic model for a single-cell, in the absence of FRH1 diffusion. **B** Stability diagram for a single cell (without FRH1 diffusion) showing the regions in the $\gamma - \alpha_0$ parameter space where epidermal (OFF), rhizoid (ON) or both states (bistable) are possible. The colormap represents the steady-state value of $R$ for each pair of parameters, starting with initial conditions with high $R$. Black lines are the same as in Fig. 3A **C** Bifurcation diagram showing the steady states of $R$ as a function of the inhibition strength $\gamma$ (and $\alpha_0 = 0.4$). For low values of $\gamma$, the system displays bistability, where cells become epidermal or rhizoid precursors depending on their initial conditions. There is a threshold $\gamma_c$ such that for $\gamma > \gamma_c$, bistability disappears and only the epidermal state is possible. However, close to the bifurcation point, a ghost remnant remains, leading to long transients in the system dynamics. **D** If the system starts close to the saddle-node bifurcation, the time it takes to reach the steady state depends on the difference $\phi \equiv \gamma - \gamma_c$, where $\gamma_c$ is the value at the bifurcation point. **E** In the ghost region, the scaling relation between the characteristic time $\tau_c$ and the proximity to the saddle-note bifurcation $\phi$ follows the power law $\tau_c \sim \phi^\beta$, with scaling exponent $\beta = -1/2$. For all panels, rest of the parameters as in Table 1.

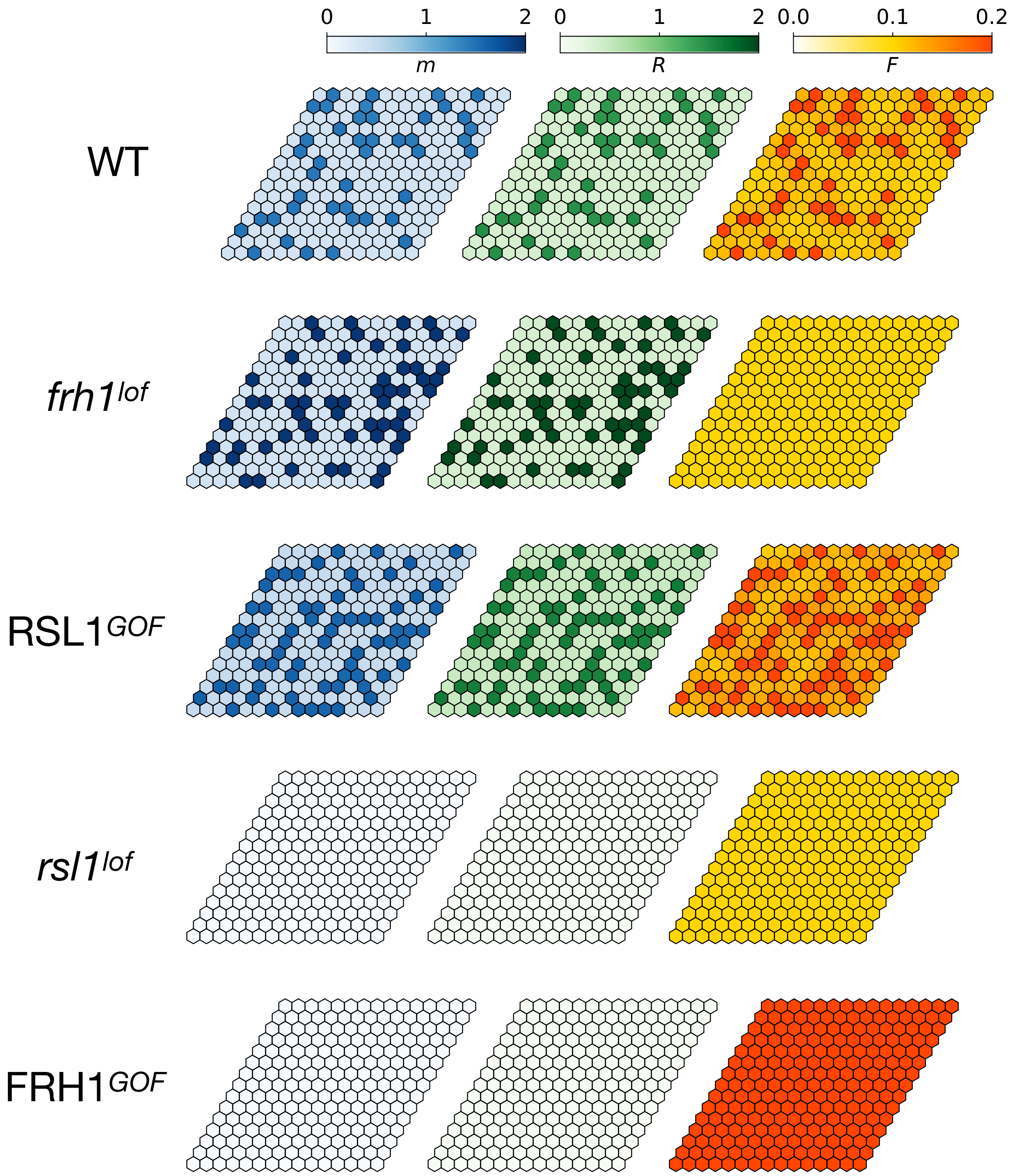


**Figure S2: Rhizoid precursor patterns in terms of each variable of the mechanistic model for WT and mutants.** Each row illustrates a typical pattern of the genotype indicated on the left. Colors indicate the concentrations of RSL1 mRNA (*m*, blue, left), RSL1 protein (*p*, gree, middle) and FRH1 miRNA (*F*, orange, right). The scale of colobars apply to all genotypes. In the WT, rhizoid precursors — cells with high values of $R$ — also have high concentration of $m$ and $F$. Because $F$ diffuses, the concentration of $F$ in epidermal cells adjacent to rhizoid precursors is intermediate. $frh1^{lof}$ and $RSL1^{GOF}$ display more rhizoid precursors, and larger and less filamentous clusters; in $rsl1^{lof}$ and $FRH1^{GOF}$ mutants no rhizoid precursors emerge. Rhizoid precursors in $frh1^{lof}$ have higher concentration of $R$ and $m$ than in the WT, but all cells have the same, low concentration of $F$. In $RSL1^{GOF}$ mutants, the pattern can be observed for all variables. For every genotype, parameters and initial conditions are as in Table 1.

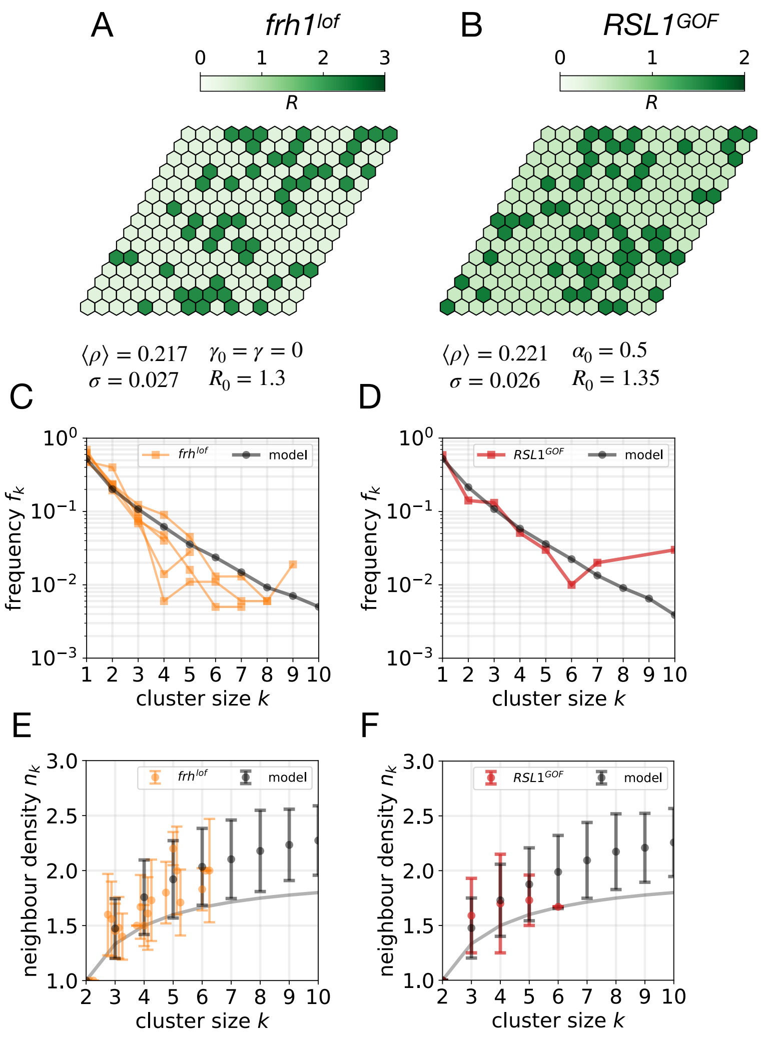


**Figure S3: Mutant phenotypes can be modelled with different parameter sets. A-B** Typical patterns of $R$ in *frh1*$^{lof}$ (A) and *RSL1*$^{GOF}$ (B) mutants for different values of the parameters and initial conditions. The corresponding values are shown below the lattices. **C-D** Cluster-size distributions for *frh1*$^{lof}$ (C) and *RSL1*$^{GOF}$ (D) mutants. **E-F** Mean neighbor density for *frh1*$^{lof}$ (E) and *RSL1*$^{GOF}$ (F) mutants. For each genotype, the values of $\langle\rho\rangle$, $\sigma$, $f_k$ and $n_k$ in the model have been computed from $N_{sim} = 1000$ simulations using lattices of size $15 \times 15$. Rest of the parameters as in Table 1.

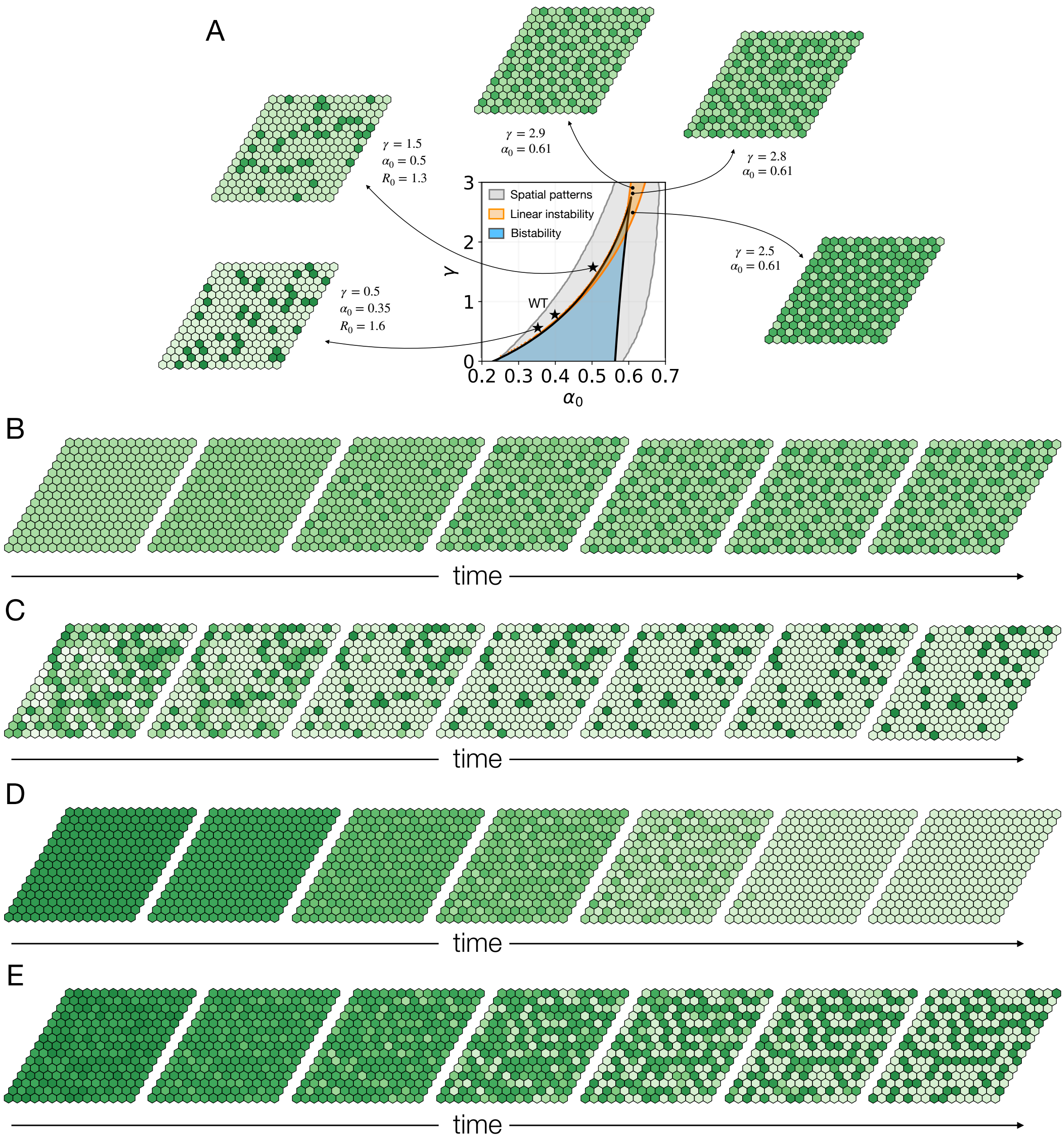


**Figure S4: Rhizoid precursor patterns in different regions of the parameter space for the mechanistic model. A** Stationary patterns of $R$ obtained for different sets of $(\gamma, \alpha_0)$ pairs. The parameter space of Fig. 3A is repeated here as a guide. Patterns that are consistent with WT gemmae only appear close to the saddle-node bifurcation (black stars; two additional cases are depicted). Different values of $R_0$ have been used to obtain such patterns. In the orange-shaded area (Turing region), periodic patterns can appear due to a diffusion-driven linear instability of a high-$R$ homogeneous state. When two homogeneous states are possible (i.e. within the bistable region; blue-shaded area), only the high-$R$ homogeneous state is linearly unstable. The spatial statistics of the patterns emerging from this region of linear instability are not consistent with WT gemmae. **B** Temporal sequence of pattern initiation in the Turing region, with parameters $\alpha_0 = 0.6$ and $\gamma = 2.6$, and initial conditions close to the homogeneous steady state, $R(t=0) = R_h(1+0.001\mathscr{U}[-1,1])$. Small differences in neighboring cells become amplified due to a linear instability of the homogeneous state, leading to propagating fronts of pattern formation. **C** Temporal sequence in a similar region to the WT, with parameters $\alpha_0 = 0.35$ and $\gamma = 0.5$, and random initial conditions ($R(t=0) = R_0\mathscr{U}[0,1]$) with $R_h = 1.6$. The resulting patterns resemble those observed in WT gemmae. **D** Temporal sequence for the same parameters as the WT (Table 1, i.e. with $\alpha_0 = 0.4$ and $\gamma = 0.75$) but for initial conditions with high-$R$ and low variability, $R(t=0) = R_h(1+V\mathscr{U}[-1,1])$, with $R_h = 1.53$ and $V = 0.01$. Even though the system starts with all cells having high $R$, it eventually collapses to the low-$R$ steady state without any rhizoid precursor. **E** Temporal sequence for the same parameters and initial conditions as in D except for higher initial variability $V = 0.1$. In this case, the higher variability allows the system to reach an inhomogeneous, spatially localized state with high density and an overrepresentation of long filamentous clusters, being distinct to that found in WT gemmae. In all panels, the remaining parameter values are shown in Table 1; initial conditions for $m$ and $F$ are zero. The color scale is the same as in Fig. 3G.

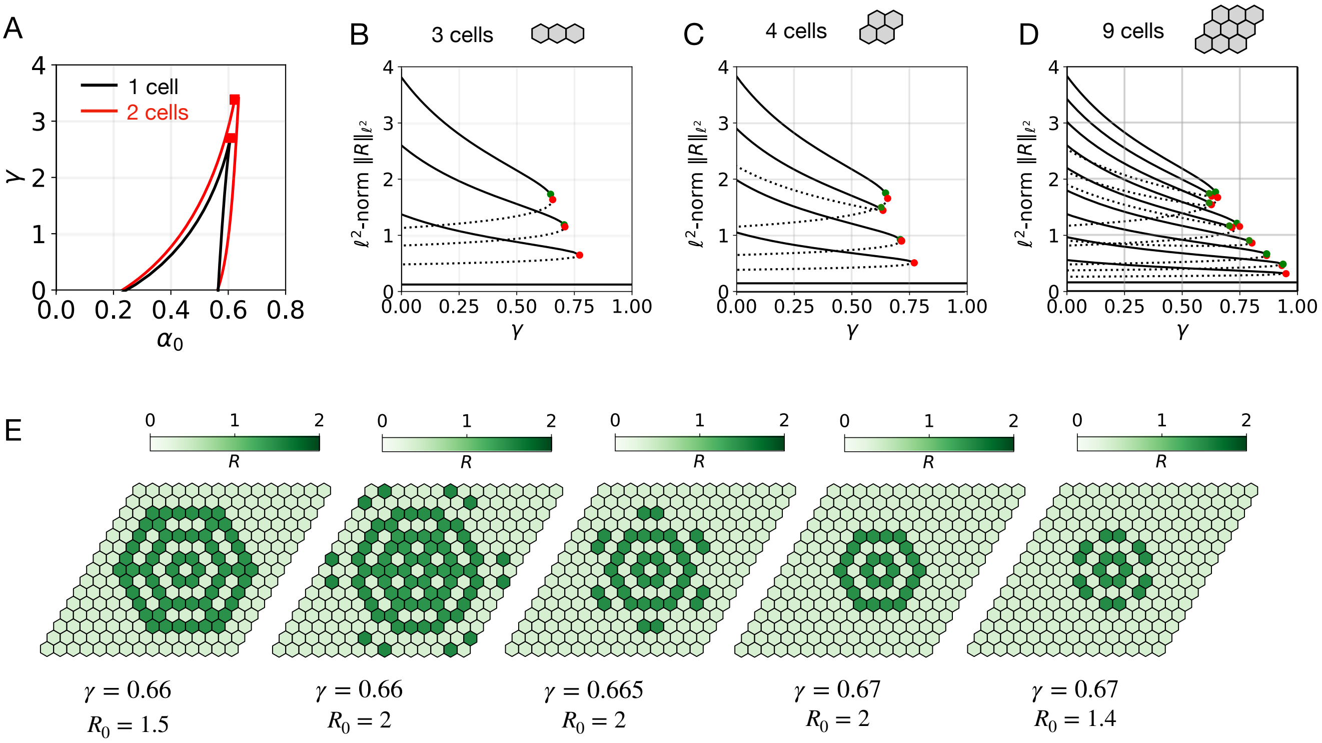


**Figure S5: Diffusion of FRH1 facilitates the bistable switch and creates patterns.** **A** Stability diagram of the mechanistic model as a function of $\gamma$ and $\alpha_0$, showing the continuation of saddle-node bifurcations for one isolated cell (black lines) and for two cells coupled by diffusion of FRH1 and isolated from any other cell (red lines). Red squares denote cusp points, where two saddle-node bifurcations collide and annihilate each other. In the parameter region enclosed by the black lines, the isolated cell can become either a rhizoid precursor or an epidermal cell, i.e. there is bistability of cell states. This region is the one depicted in blue in Fig. 3A. In the same region, cells in a lattice interacting via the diffusible FRH1 can become all rhizoid precursors or all epidermal. Outside this region, an isolated cell can only reach one fate — there is no bistability. The parameter region enclosed by the red lines denotes the bistable region for two coupled cells. Outside this region, the two cells can only reach a single fate. **B-D** Bifurcation diagrams showing the $\ell^2$-norm $||R||_{\ell^2}$ as a function of $\gamma$, for 3 (B), 4 (C) and 9 (D) cells interacting via the diffusion of FRH1, with periodic boundary conditions. In panels B-E, $\alpha_0 = 0.4$. Solid and dotted lines denote stable and unstable steady states, respectively. The solid line with largest values of $||R||_{\ell^2}$ represents the state where all cells are rhizoid precursors; the horizontal solid line with lowest value of $||R||_{\ell^2}$ corresponds to all cells being epidermal. Regardless of the cell number, these two lines are the same in all panels; however, as the number of interacting cells increases, more intermediate states appear. Each of these states correspond to inhomogeneous patterns with a spatial distribution of rhizoid precursors and epidermal cells. The higher the number of cells, the larger the inhibition strength $\gamma$ has to be to completely inhibit rhizoid precursor specification (i.e. so the only stable state is all cells being epidermal). Red circles denote saddle-node bifurcations, while green circles denote pitchfork bifurcations. **E** Localized patterns obtained with different values of $\gamma$, for initial conditions consisting of all cells with $R_0 = 0$, except the cell in the middle, which has $R_0$ equal to the values shown below the lattices. In all panels, the rest of the parameters are shown in Table 1, and initial conditions for $m$ and $F$ are set to zero.

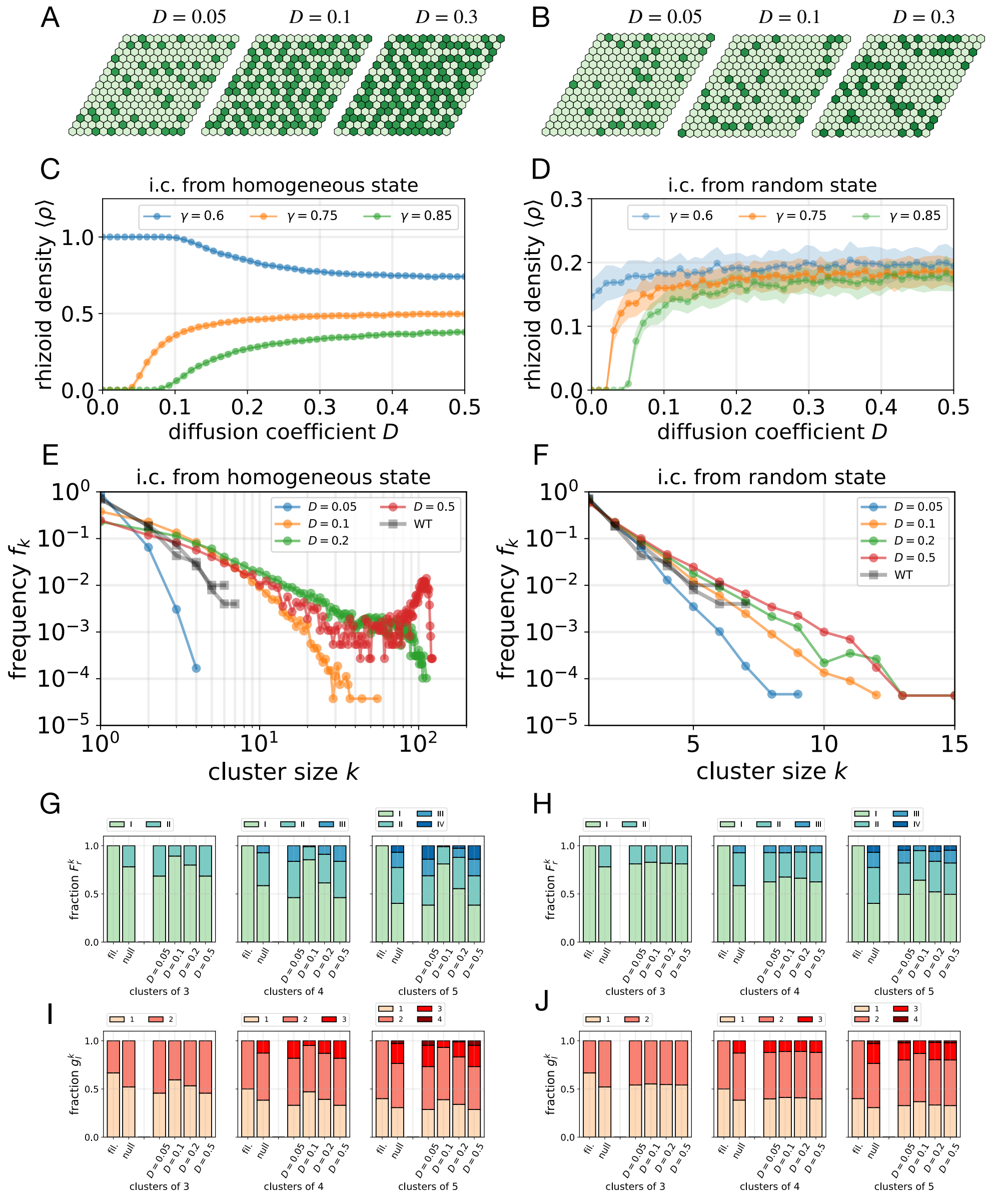


**Figure S6: Effect of the diffusion coefficient and the initial conditions.** Results on the left side (A,C,E,G and I) correspond to initial conditions with all cells in a high-$R$ state and some variability ($R(t=0) = R_h(1+V\mathscr{U}[-1,1])$ with $R_h \simeq 1.4$ and $V = 0.1$). Results on the right side (B,D,F,H and J) correspond to the same parameter values as in the left side but for strong random initial conditions, with only some cells initially having high $R$ ($R(t=0) = R_0\mathscr{U}[0,1]$ with $R_0 = 1.53$). **A, B** Examples of stationary rhizoid precursor patterns for different values of the diffusion coefficient $D$. In all cases, $\gamma = 0.75$. **C, D** Stationary density of rhizoid precursors for different diffusion coefficients $D$ and different $\gamma$. Points denote averages over $N = 41$ simulations, and shaded regions $\pm$ standard deviations. **E, F** Stationary cluster-size distributions for different values of $D$. Each curve is obtained from $N = 1000$ simulations. The scale and limits of the $x$ axes in E and F are different for clarity. In F, the frequency $f_k$ is much less sensitive to the diffusion coefficient $D$ than in E. **G, H** $F_r^k$ for different values of $D$. Bars labeled by *fil.* and *null* show the results computed for filamentous clusters and for the null model ($p = 0.15$, $N = 1000$), respectively. **I, J** $g_l^k$ for different values of $D$, using the same simulations as in G and H, respectively. Panels G,I and H,J correspond to the same simulations in E and F, respectively. Results in G,I and H,J show that an intermediate value of $D$ is needed to generate lateral inhibition. If $D$ is too large, inhibition in no longer local, and the statistics of clusters behave as in the null model. In all panels, the rest of the parameters is shown in Table 1. Simulations are performed on lattices of $15 \times 15$ cells.

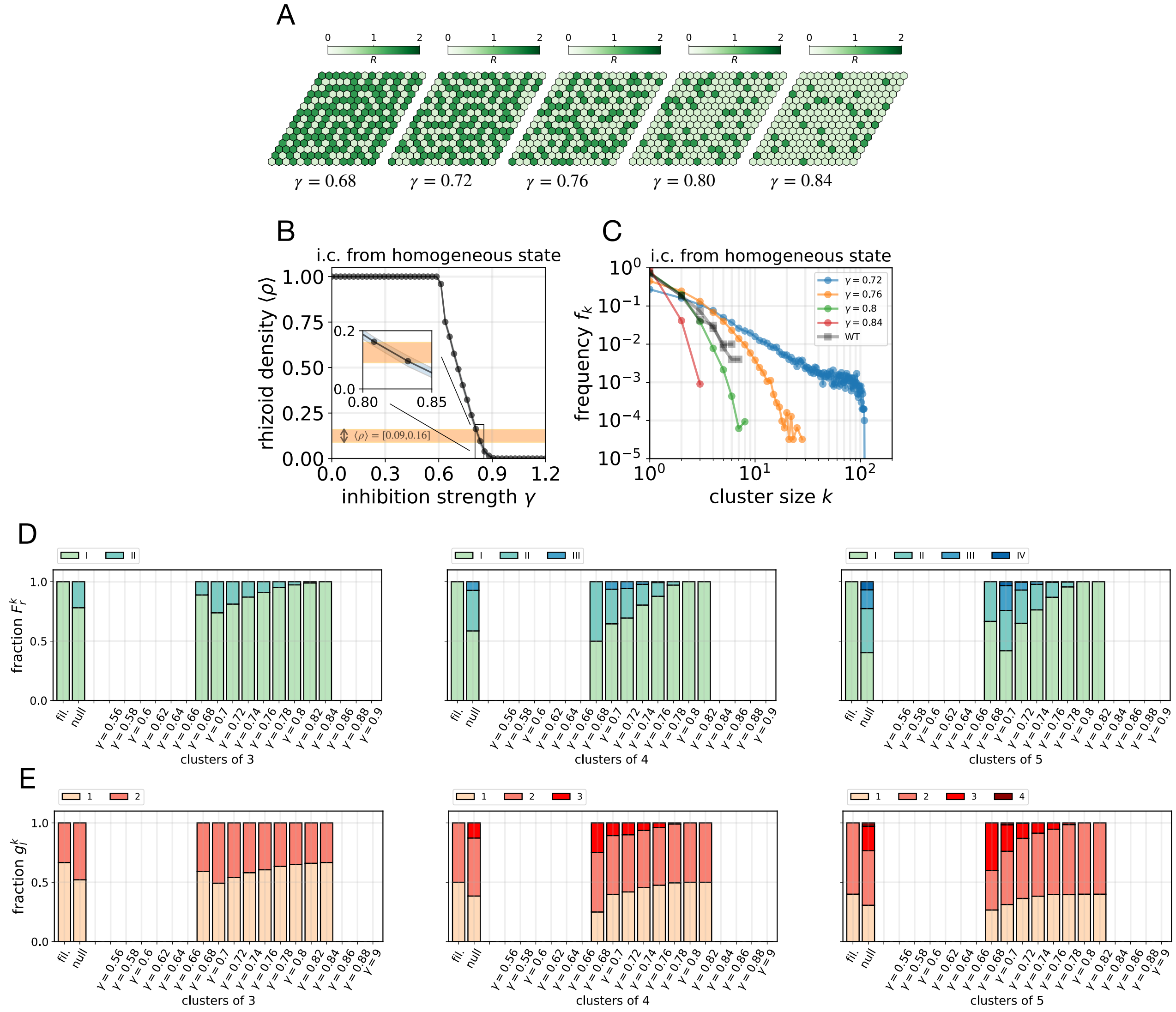


**Figure S7: Results of the mechanistic model when all cells start with high levels of RSL1.** Results of the mechanistic model for parameter values as in Fig. 4 but for initial conditions with all cells having high $R$, and some variability between cells ($R(t=0) = R_h(1+V\mathscr{U}[-1,1])$, where $R_h$ is the value in the high-$R$ homogeneous state and $V = 0.1$). **A** Example of stationary patterns for different inhibition strengths $\gamma$. The random initial conditions are different for each value of $\gamma$, but with the same variability $V$. **B** Stationary average rhizoid density for different inhibition strengths. The homogeneous state with all cells being rhizoid precursors (i.e. average density equal to 1) is stable to small perturbations up to a threshold of the inhibition strength $\gamma \approx 0.6$. For higher $\gamma$ there is a sharp decrease in rhizoid density. The orange strip represents the region where the average density of rhizoid precursors is consistent with experimental observations, $\langle \rho \rangle \in (0.09, 0.16)$. Each point represents the average over $N = 41$ simulations. The blue shaded area (only visible in the inset) represents $\pm$ the standard deviation. **C** Stationary cluster-size distribution for different values of $\gamma$. Each distribution is computed from $N = 1000$ simulations. Initial conditions as in A and B. Grey data correspond to the WT distributions described in [1]. For these high-$R$ initial conditions it is not possible to find a value of $\gamma$ that reproduces both the average density and $f_k$ of WT gemmae. The values of $f_k$ for these initial conditions are much more sensitive to $\gamma$ than for high initial variability (Fig. 4E). **D** and **E** show the values of $F_r^k$ and $g_l^k$, respectively, obtained from the the same simulations ($N = 1000$) for clusters of size $k = 3$ (left), $k = 4$ (middle) and $k = 5$ (right). Higher $\gamma$ leads to more filamentous clusters (Type I) independently of the cluster size. The absence of bars for $\gamma \leq 0.66$ reflects the fact that all clusters are very large ($k > 5$), as inhibition is to low to produce small clusters, and therefore no clusters of size $k = 3, 4, 5$ appear. For $\gamma \geq 0.86$, inhibition is too strong to produce rhizoid precursor clusters of size 3 or more, and therefore there are no statistics. Panels D and E can be compared to Fig. 4 B,C, which show the corresponding results when the initial conditions have low average $R$ and high variability, such that only some cells initially have high $R$. The values of $F_r^k$ and $g_l^k$ shown in Fig. 4 B,C corresponding to the null model and to filamentous clusters are also plotted here to facilitate comparison. For all panels, rest of the parameters are as in Table 1 and simulations are performed on lattices of $15 \times 15$ cells.

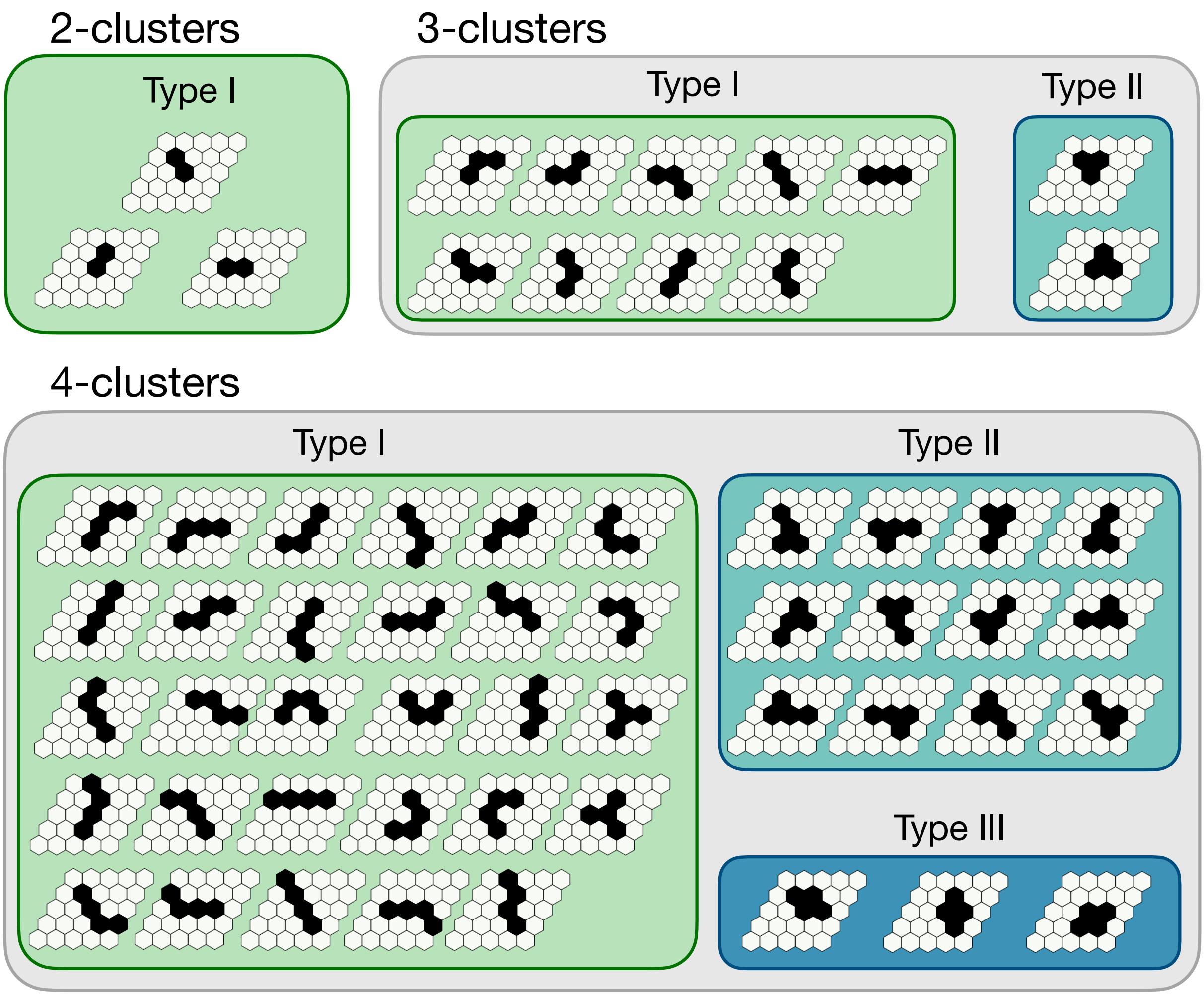


**Figure S8: All possible spatial configurations and cluster types for clusters of $k = 2,3,4$ rhizoid precursors.** In a regular hexagonal lattice, rhizoid clusters can appear with a number of spatial configurations that depends on the cluster size $k$. These configurations can be classified into cluster types. For every size $k$, Type I corresponds to filamentous configurations. Two features are common for spatial configurations of the same size and type: 1) the number of rhizoid precursors with $l$ adjacent rhizoids (and consequently, the average number of adjacent rhizoid precursors $n_k^r$) and 2) the total number of epidermal cells in contact with the cluster. For instance, clusters of size $k = 3$ and type I have only one rhizoid which is adjacent to $l = 2$ rhizoids, and two rhizoids which are adjacent to only $l = 1$ rhizoid. Therefore $n_3^l = \frac{1\times2+2\times1}{3} = 4/3$, and with 10 epidermal cells surrounding the clusters. The type of mechanism involved in rhizoid cluster formation (e.g. lateral inhibition) will have an effect on the frequencies of each cluster size and type.

### Supplementary Tables

| Size $k$ | Type I | Type II | Type III | TOTAL |
|---|---|---|---|---|
| **1** | 419 | – | – | 419 |
| **2** | 120 | – | – | 120 |
| **3** | 32 | 5 | – | 37 |
| **4** | 17 | 4 | 0 | 21 |
| **5** | 5 | 1 | 0 | 6 |
| **6** | 3 | 1 | 0 | 4 |
| **7** | 0 | 0 | 0 | 0 |
| **8** | 1 | 0 | 0 | 1 |
| **9** | 0 | 0 | 1 | 1 |

**Table S2:** Experimental results on the statistics of rhizoid precursor clusters ($N = 43$ gemmae).

| Size $k$ | Type I | Type II | Type III | Type IV | TOTAL |
|---|---|---|---|---|---|
| **1** | 14537 | – | – | – | 14537 |
| **2** | 4691 | – | – | – | 4691 |
| **3** | 1462 | 306 | – | – | 1768 |
| **4** | 525 | 179 | 42 | – | 746 |
| **5** | 149 | 100 | 10 | 1 | 260 |
| **6** | 63 | 39 | 7 | 1 | 110 |
| **7** | 28 | 23 | 4 | 0 | 55 |
| **8** | 9 | 4 | 1 | 1 | 15 |
| **9** | 3 | 3 | 0 | 0 | 6 |
| **10** | 0 | 1 | 1 | 0 | 2 |

**Table S3:** Statistics of rhizoid precursor clusters for $N = 1000$ simulations of the mechanistic model (Fig. 5)

| Size $k$ | Type I | Type II | Type III | Type IV | Type V | Type VI | Type VII | Type VIII | TOTAL |
|---|---|---|---|---|---|---|---|---|---|
| **1** | 13989 | – | – | – | – | – | – | – | 13989 |
| **2** | 4255 | – | – | – | – | – | – | – | 4255 |
| **3** | 1304 | 341 | – | – | – | – | – | – | 1645 |
| **4** | 383 | 221 | 56 | – | – | – | – | – | 660 |
| **5** | 130 | 118 | 29 | 15 | – | – | – | – | 292 |
| **6** | 44 | 56 | 30 | 9 | 5 | – | – | – | 144 |
| **7** | 21 | 20 | 15 | 6 | 2 | 1 | – | – | 65 |
| **8** | 5 | 11 | 16 | 6 | 5 | 2 | 0 | – | 45 |
| **9** | 9 | 6 | 1 | 2 | 2 | 1 | 0 | 0 | 21 |
| **10** | 3 | 3 | 1 | 1 | 1 | 0 | 0 | 0 | 9 |
| **11** | 2 | 0 | 0 | 1 | 1 | 0 | 0 | 0 | 4 |
| **12** | 0 | 0 | 0 | 1 | 0 | 1 | 0 | 1 | 3 |
| **13** | 0 | 0 | 1 | 0 | 1 | 1 | 0 | 0 | 3 |
| **17** | 0 | 0 | 0 | 1 | 0 | 0 | 0 | 0 | 1 |

**Table S4:** Statistics of rhizoid precursor clusters for $N = 1000$ simulations of the null model with $p = 0.15$ (Fig. 1)

| | Exp. data | Mech. model | null model |
|---|---|---|---|
| **Type I** | 33 | 1462 | 1304 |
| **Type II** | 4 | 306 | 341 |
| *p*-value (Fisher) | | 0.3821 | 0.2137 |
| *p*-value ($\chi^2$) | | | 0.01226 |

**Table S5:** Contingency tables of the total numbers of clusters of size $k = 3$ of each type (I,II) from $N_{gemmae} = 43$ WT gemmae (Exp. data), $N = 1000$ simulations of the mechanistic model in WT conditions (Mech. model) and $N = 1000$ simulations of the null model with $p = 0.15$ (null model). The *p*-value (Fisher) denoted for each model column is the result of Fisher test on the contingency table for that model and the Exp. data column. The *p*-value ($\chi^2$) in the last column is the result from $\chi^2$-test on the contingency table from the results of the Mech. model and null model. These data are represented in Fig. 5F.

| | Exp. data | Mech. model | null model |
|---|---|---|---|
| **Type I** | 26 | 777 | 597 |
| **Type II** | 6 | 349 | 435 |
| **Type >II** | 1 | 68 | 216 |
| *p*-value (Fisher) | | 0.2949 | 0.001361 |
| *p*-value ($\chi^2$) | | | <2.2e-16 |

**Table S6:** Contingency tables of the total numbers of clusters of size $k \geq 4$ of each type (I,II) from $N_{gemmae} = 43$ WT gemmae (Exp. data), $N = 1000$ simulations of the mechanistic model in WT conditions (Mech. model) and $N = 1000$ simulations of the null model with $p = 0.15$ (null model). The *p*-value (Fisher) denoted for each model column is the result of Fisher test on the contingency table for that model and the Exp. data column. The *p*-value ($\chi^2$) in the last column is the result from $\chi^2$-test on the contingency table from the results of the Mech. model and null model. These data are represented in Fig. 5G.

| | Exp. data | Mech. model | null model |
|---|---|---|---|
| $l = 1$ | 66 | 2924 | 2608 |
| $l = 2$ | 45 | 2380 | 2327 |
| *p*-value (Fisher) | | 0.3865 | 0.1789 |
| *p*-value ($\chi^2$) | | | 0.02176 |

**Table S7:** Contingency tables of the total number of cells with $l = 1,2$ adjacent rhizoid precursors in clusters of size $k = 3$ from $N_{gemmae} = 43$ WT gemmae (Exp. data), $N = 1000$ simulations of the mechanistic model in WT conditions (Mech. model) and $N = 1000$ simulations of the null model with $p = 0.15$ (null model). These results can be obtained from Table S5 (see Supp. Info Text). The *p*-value (Fisher) denoted for each model column is the result of Fisher test on the contingency table for that model and the Exp. data column. The *p*-value ($\chi^2$) in the last column is the result from $\chi^2$-test on the contingency table from the results of the Mech. model and null model. These data are represented in Fig. 5G.

| | Exp. data | Mech. model | null model |
|---|---|---|---|
| $l = 1$ | 62 | 2149 | 2013 |
| $l = 2$ | 82 | 2731 | 2855 |
| $l > 2$ | 11 | 643 | 1326 |
| *p*-value (Fisher) | | 0.2103 | 1.013e-05 |
| *p*-value ($\chi^2$) | | | <2.2e-16 |

**Table S8:** Contingency tables of the total number of cells with $l = 1,2,> 2$ adjacent rhizoid precursors in clusters of size $k > 3$ from $N_{gemmae} = 43$ WT gemmae (Exp. data), $N = 1000$ simulations of the mechanistic model in WT conditions (Mech. model) and $N = 1000$ simulations of the null model with $p = 0.15$ (null model). The *p*-value (Fisher) denoted for each model column is the result of Fisher test on the contingency table for that model and the Exp. data column. The *p*-value ($\chi^2$) in the last column is the result from $\chi^2$-test on the contingency table from the results of the Mech. model and null model. These data are represented in Fig. 5G.

### References

[1] Anna Thamm, Timothy E. Saunders, and Liam Dolan. MpFEW RHIZOIDS1 miRNA-Mediated Lateral Inhibition Controls Rhizoid Cell Patterning in Marchantia polymorpha. *Current Biology*, 30:1905–1915, 2020.
